# Maintaining structural and functional homeostasis of the *Drosophila* respiratory epithelia requires stress-modulated JAK/STAT activity

**DOI:** 10.1101/2020.06.19.160929

**Authors:** Xiao Niu, Christine Fink, Kimberley Kallsen, Leizhi Shi, Viktoria Mincheva, Sören Franzenburg, Ruben Prange, Iris Bruchhaus, Judith Bossen, Holger Heine, Thomas Roeder

## Abstract

Signaling mediated by the Janus kinase (JAK)/Signal Transducer and Activator of Transcription (STAT) pathway is critical for maintaining cellular and functional homeostasis in the lung. Thus, chronically activated JAK/STAT signaling is causally associated with lung diseases such as lung cancer, asthma, and chronic obstructive pulmonary disease. To elucidate the molecular processes that transform increased JAK/STAT signaling in airway epithelial cells into the known pathological states, we used a highly simplified model system, the fruit fly *Drosophila melanogaster*. Here, the JAK/STAT pathway is permanently active in almost all airway cells and responds to airborne stressors with increased activity. Silencing of this signaling pathway in epithelial cells resulted in apoptosis. Since the above-mentioned lung diseases are commonly associated with increased JAK/STAT signaling, we assessed this by its ectopic activation in the respiratory epithelium of *Drosophila*. This intervention triggered cell-autonomous structural changes in epithelial cells. These structural changes included phenotypes associated with asthma, namely, thickening of the epithelium, substantial narrowing of the air-conducting space, and impairment of the secretory epicuticular structure of the tracheae. Pharmacological manipulation of JAK/STAT signaling reversed this pathological phenotype. Transcriptomic analyses revealed that several biological processes were affected, which is consistent with the impairment of junction protein trafficking also observed in this study. These results indicate that balanced JAK/STAT signaling is essential for the functionality of the respiratory epithelium and, by extension, the entire organ. In contrast, chronic overactivation of this signaling leads to massive structural changes that are closely associated with pathologies typical of chronic inflammatory lung diseases.

**Highlights:**

1. JAK/STAT signaling is active in the entire *Drosophila* respiratory system in all developmental stages.
2. The signaling pathway is indispensable for the survival of the tracheal cell.
3. Overactivation of the signaling has significant effects on tracheal development and also displays a human disease-associated phenotype in *Drosophila* trachea.

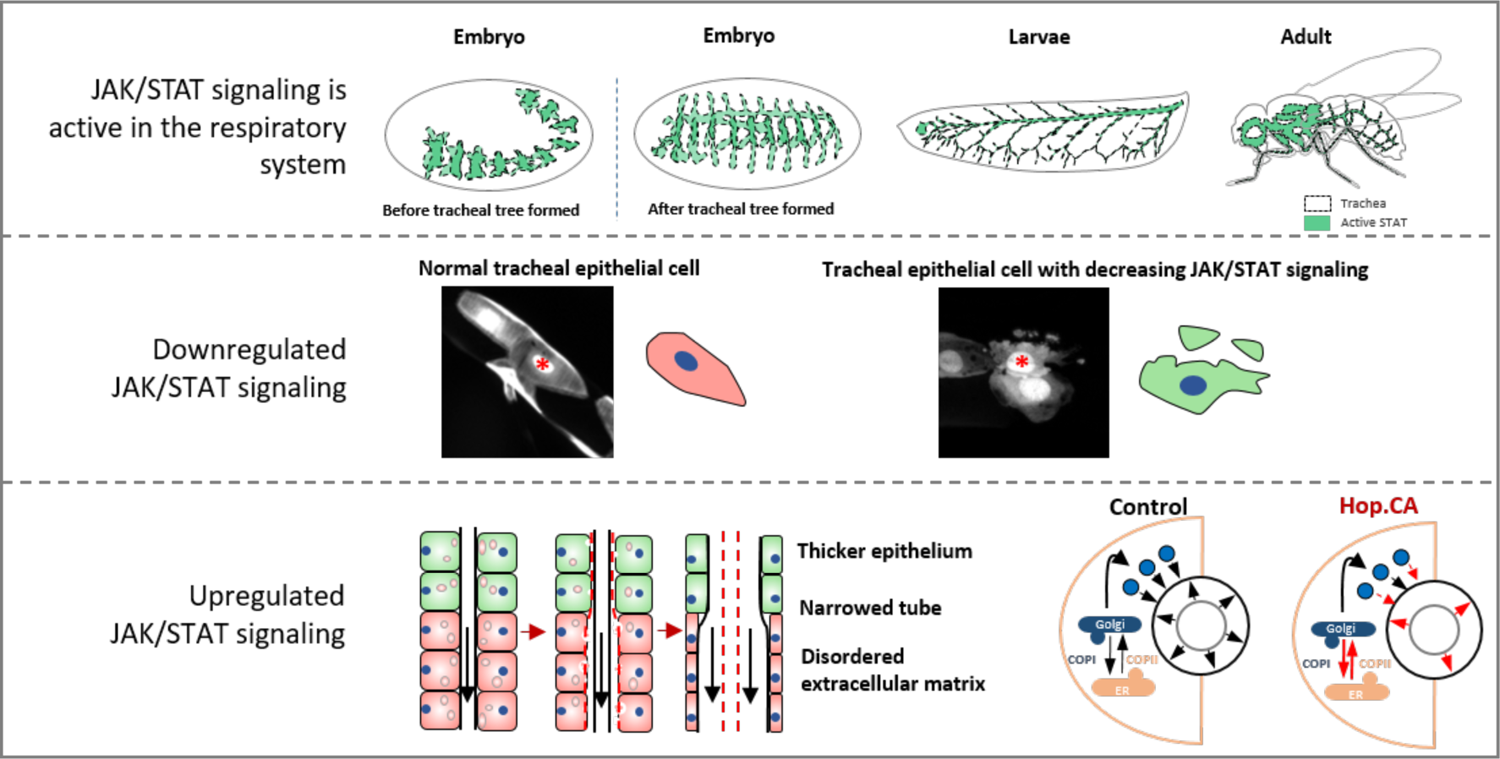

## Introduction

The Janus kinase (JAK)/signal transducer and activator of transcription (STAT) signaling system is of central importance for several critical physiological processes such as development, tissue homeostasis, and immune responses (Philips et al., 2022; Rawlings et al., 2004). Signal transduction via this pathway is straightforward and allows environmental factors to directly influence transcriptional activity, linking key biological processes to the environment (Kiu and Nicholson, 2012). Deregulation of this pathway is linked to numerous human diseases, with cancer and inflammatory pathologies being particularly prominent (Hu et al., 2021; Milara et al., 2018). JAK/STAT signaling acts downstream of a plethora of cytokines that transmit immune-related information, acting as a central integration hub in almost all cells of the lung (Yew-Booth et al., 2015)(Villarino et al., 2017). Signaling via this pathway is essential during organ development and for maintaining tissue and immune homeostasis of the fully differentiated lung. Here, JAK/STAT signaling is required to cope with stressors, infection, and damage (Jin et al., 2018; Kida et al., 2008; Major et al., 2020; Makris et al., 2017; Tadokoro et al., 2014). Given the importance of JAK/STAT signaling in the lung, it is predictable that decreased or increased signaling via this pathway leads to pathologies. Whereas reduced JAK/STAT signaling corresponds with impaired repair capacities (Tadokoro et al., 2014), increased JAK/STAT signaling is associated with a plethora of chronic lung diseases including asthma, chronic obstructive pulmonary disease (COPD), idiopathic pulmonary fibrosis (IPF), and lung cancer (Dutta et al., 2014; Georas et al., 2021; Milara *et al*., 2018; Yew-Booth *et al*., 2015). Despite its simple general organization, the JAK/STAT signaling pathway in vertebrates shows multiple redundancies, parallels, and convergences; a situation that is further complicated since these signaling pathways may act differently in different cell types of the same organ (Hu *et al*., 2021; Morris et al., 2018; Villarino *et al*., 2017). Therefore, models with a much simpler JAK/STAT signaling pathway and a less complex cellular composition in the airways should help to elucidate the effects of deregulated JAK/STAT signaling, especially in airway epithelial cells. *Drosophila melanogaster* can be used for this task, as only one receptor (Domeless), one JAK kinase (Hopscotch), and one STAT transcription factor (STAT92E) are present (Arbouzova and Zeidler, 2006; Zeidler and Bausek, 2013). This low level of redundancy offers the unique advantage to study the generic relevance of JAK/STAT signaling. Furthermore, the use of *Drosophila* allows focusing exclusively on the airway epithelium, a resident cell population playing a central role in the orchestration of organ homeostasis, but also for developing chronic pathologies. In the tracheal system of *Drosophila*, the lung’s functional equivalent, there is so far only information on the relevance of JAK/STAT signaling during embryonic development and the formation of the adult tracheal system (Brown et al., 2001; Perrimon and Mahowald, 1986; Powers and Srivastava, 2019). A lack of JAK/STAT signaling during very early phases of tracheal development impairs tracheal development including cell movement and elongation, as well as invagination processes that lead to tube formation (Brown et al., 2001; Isaac and Andrew, 1996). During the development of the vertebrate lung, very similar processes are operative (Nogueira-Silva et al., 2006; Piairo et al., 2018). In recent years, *Drosophila* served as a model for numerous human diseases (Pandey and Nichols, 2011) including those of the lung (Bossen et al., 2021; Levine and Cagan, 2016; Prange et al., 2018; Roeder et al., 2009; Roeder et al., 2012).

The current study aimed to elucidate the general importance of JAK/STAT signaling in the airways and understand how chronic deregulation of this pathway in airway epithelial cells results in pathological outcomes. Applying a tailored *Drosophila* model, we could not only show that JAK/STAT signaling is required to prevent apoptosis of airway epithelial cells but that a wide variety of stressors increased signaling via this pathway significantly. Ectopic activation on the other hand, leads to massive structural changes that drastically limit airway epithelial functionality. Using this model, we demonstrated that balanced JAK/STAT signaling in airway epithelia is imperative to prevent the development of pathology and that pharmacological intervention at precisely this point is excellent for addressing it.

## Results

Activation of the JAK/STAT pathway occurs in a wide variety of organs and can easily and reliably be visualized in *Drosophila* using transcriptional reporters. Here, the activity of the pathway (Fig. 1A) is visualized by a STAT92E-promoter-dependent GFP-based reporter approach (Bach et al., 2007). It was already known that the JAK/STAT pathway is strongly activated in the trachea during embryogenesis and it could be assumed to be important for tracheal development (Bach et al., 2007). However, the exact pattern of its activation is still unclear. To fill this gap of knowledge, we analyzed the *STAT92E*-reporter activity concurrently with *btl>LacZ.nls* (Shiga et al., 1996), which specifically labels the trachea. Activation of the *STAT92E-GFP* reporter was pronounced in a central region in the trachea, mainly in the transverse connective area (Fig. 1B–D). To evaluate if the JAK/STAT pathway also operates in a functional airway epithelium under control conditions, we used the *STAT92E-GFP* reporter line to monitor pathway activity in the larval tracheal system (Fig. 1E). JAK/STAT activity was relatively low in the posterior zone of the trachea (Fig. 1F), the tenth tracheal metamere (Tr10), where cells progressively undergo apoptosis in response to trachea metamorphosis (Bosch et al., 2015; Chen and Krasnow, 2014). However, JAK/STAT activity was much stronger in regions such as Tr2, the spiracular branch (SB), and the dorsal branch (DB) than in nearby areas (Fig. 1G and H). In these areas with high JAK/STAT activity, cells re-enter the cell cycle at this developmental stage (Guha et al., 2008; Sato et al., 2008; Weaver and Krasnow, 2008). We next evaluated if JAK/STAT activity is observed in airway epithelial cells of adults. The same approach detected substantial reporter activity in airway epithelial cells of fully developed adults (Fig. 1I). The JAK/STAT pathway was active throughout the entire tracheal system. In addition, we evaluated the expression of the three *upds* in the larval trachea. Experiments using the corresponding enhancer-*Gal4* lines demonstrated that especially *upd2* was highly expressed in the tracheal system (Fig. S1).

**Figure 1:**
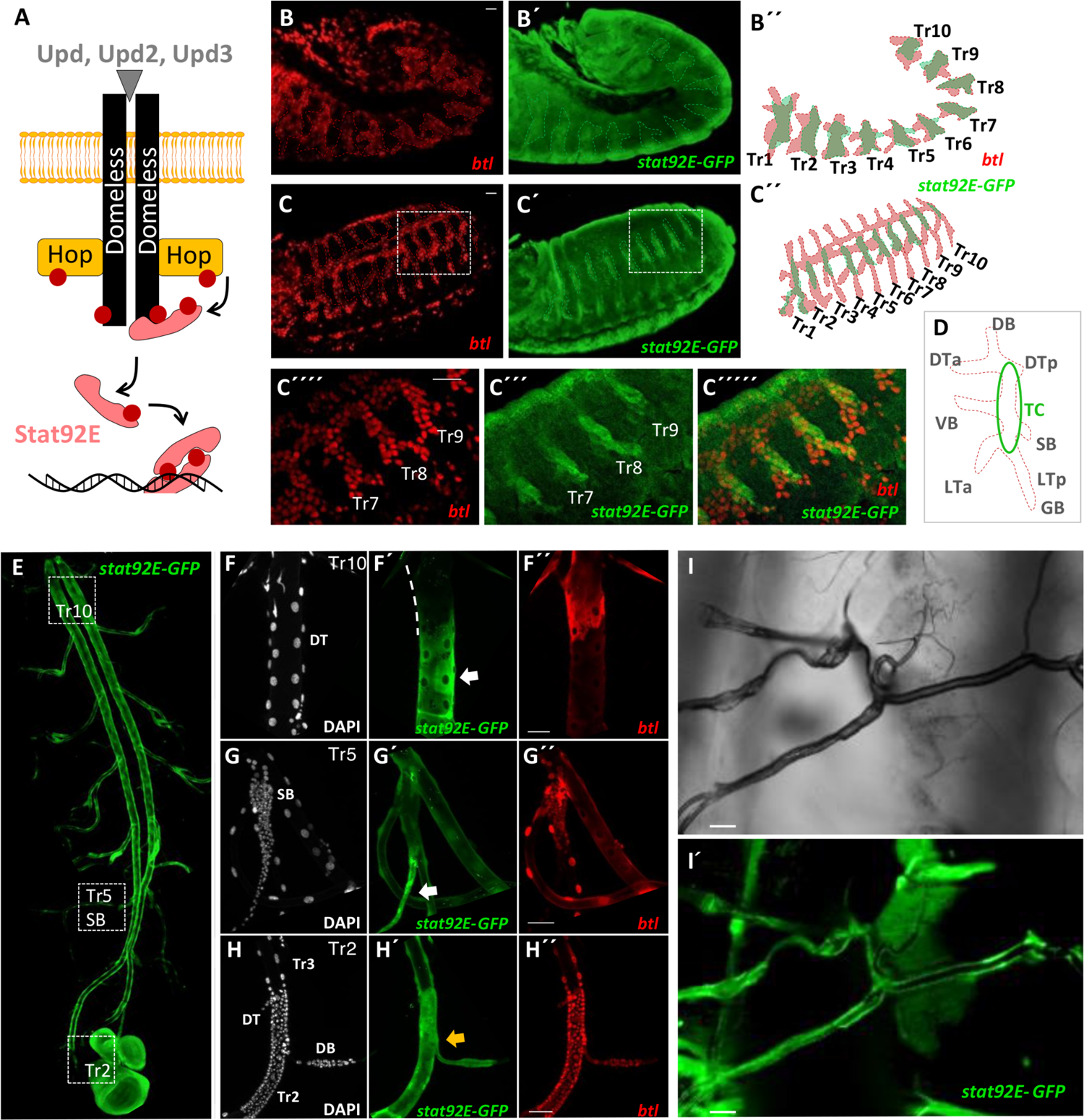
The activation of JAK/STAT signaling pathway in the respiratory system of *Drosophila*. (A) shows a model of JAK/STAT signaling pathway in *Drosophila* comprising all major components. Three ligands (Upd, Upd2 and Upd3) bind to a single receptor, Domeless (Dome), which transmits this information via a single JAK (Hopscotch (Hop)) to a single STAT transcription factor (STAT92E). The activation of JAK/STAT signaling through all developmental stages was detected by using a *STAT92E-GFP* reporter (*btl-Gal4*, *STAT92E*-*GFP*; *UAS-LacZ.nls*). (B-D) Fluorescence micrographs of embryonic trachea stained for GFP (green, JAK/STAT pathway activated zones), and beta-galactosidase (red, tracheal metamere (Tr)), respectively. The activation of JAK/STAT pathway in the terminal regions is weaker than that in the central regions before (B) and after the fuse of tracheal invaginations of all segments (C). (D) shows the activation pattern of JAK/STAT signaling in the embryonic trachea metamere. The stronger signal in the central region mainly belongs to TC (green circle). DB, dorsal branch; DTa, dorsal trunk anterior; DTp, dorsal trunk posterior; VB, visceral branch; LTa, lateral trunk anterior; LTp, lateral trunk posterior; and GB, ganglionic branch; SB, spiracular branch; transverse connective, TC. The stronger signal in the central regions mainly belongs to TC. Scale bar: 20 µm. (E-H) Fluorescence micrographs of larval trachea. JAK/STAT activity is present in most parts of the trachea (E). The region in the posterior region of the trachea (dorsal trunk (DT) of Tr10, white dash line in F) showed a decreased activity of JAK/STAT signaling compared to other tracheal epithelial cells. Scale bar: 50 µm. Conversely, some regions like SB (G, marked by white arrow) and Tr2, DB (H, marked by white arrow) showed an increased activity of JAK/STAT signaling compared to the tracheal epithelial cells. Scale bar: 50 µm. (I) Fluorescence micrographs of adult trachea. JAK/STAT signaling was induced in the trachea of the adult.

### The JAK/STAT pathway responds to stressful stimuli

To investigate whether external, stress-associated stimuli can affect JAK/STAT signaling in airway epithelia, we exposed the just-described reporter lines to these stimuli. We focused on airborne stressors and exposed both larvae and adults to them. Here, the administration of cold air (cold), hypoxia, and cigarette smoke (CSE) boosted the activity of the JAK/STAT pathway in airway epithelial cells (Figure 2A and 2C) of both larval and adult animals. A quantitative evaluation of the induced fluorescent signals showed that these increases were statistically significant (Fig. 2C).

**Figure 2:**
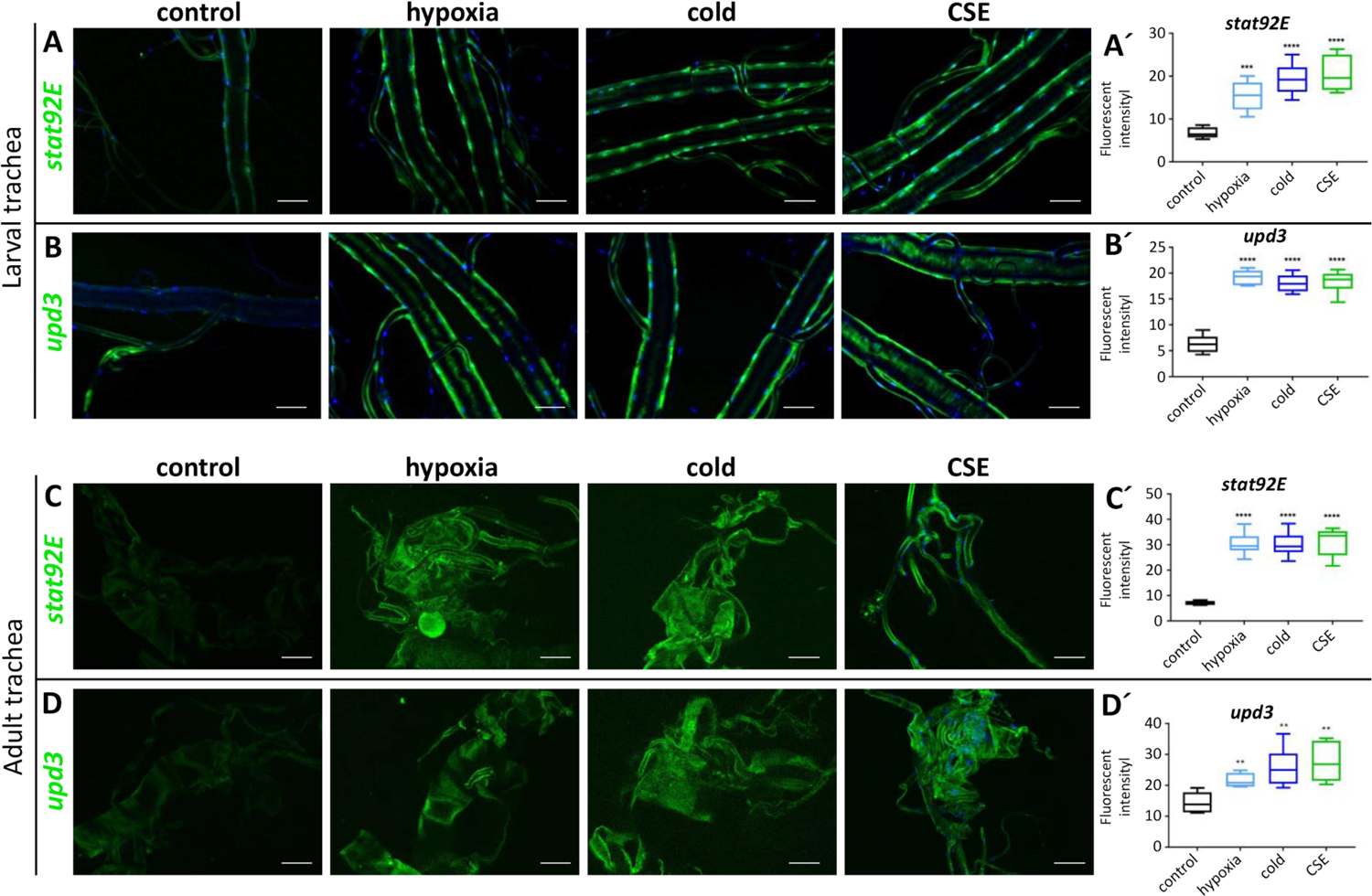
JAK/STAT signaling pathway in trachea was activated by external stimuli. (A and C) Fluorescence micrographs of the trachea of *STAT92E*-*GFP* larvae (A) and adult (C) under normal conditions and external stimuli, including hypoxia, cold air (cold), cigarette smoke exposure (CSE). The external stimuli induced activity of STAT92E could be observed and was quantified through fluorescent intensity (A’ and C’). (B and D) Fluorescence micrographs of the trachea of *upd3*-*Gal4*; *UAS-GFP* larvae (B) and adult (D) that were exposed to normal condition and external stimuli. The external stimuli induced expression of *upd3* could be observed and was quantified through fluorescent intensity (B’ and D’). ** *p* < 0.01, *** *p* < 0.001, **** *p* < 0.0001 by Student’s t-test. Scale bar: 100 µm.

Then we expanded the analysis of the effects of airborne stressors to include the ligands of the signaling system. We found that one of the three ligands, *upd*, did not show any expression independent of the situation (data not shown). In contrast, the expression of *upd2* and *upd3* were induced by external stimuli, for *upd2* only by CSE, for *upd3* by CSE and hypoxia (Fig. S2). The activity of the *STAT92E-GFP* reporter was induced exactly in the region where also the ligand showed enhanced expression. For further analyses, we focused on the most responsive *upd* gene, namely *upd3*. Expression of *upd3* was low in the tracheal cells of larvae or adult animals that lived under control conditions but could easily be observed in all these tracheal cells when the animals were exposed to external stimuli (Fig. 2B and 2D).

### JAK/STAT signaling is essential for the survival of airway epithelial cells

To evaluate the relevance of the JAK/STAT pathway in the *Drosophila* respiratory system, we blocked its activity in the trachea by driving the expression of a dominant-negative isoform of Domeless (*Dome.DN*) using the trachea-specific *btl*-*Gal4* driver. As a result, some animals developed to the larval stage (Fig. 3A), but all died before reaching the pupal stage. Microscopic analysis of surviving larvae showed that the dorsal trunk (DT) was absent (Fig. 3B’). Time-lapse imaging of embryos experiencing trachea-specific *Dome.DN* expression revealed that all tracheal segments were fused, however, the DT subsequently separated (Fig. 3C). STAT92E-RNAi was used to confirm this kind of effects on trachea. A detailed analysis of the effects caused by constitutive *Dome.DN* expression was performed using a mosaic approach (Wu et al., 2006). Here, driving expression in a mosaic fashion by *vvl-FLP*, *CoinFLP-Gal4* changed the morphology of the affected cells. These cells appear to undergo apoptosis (Fig. 3D). Apoptosis of these affected cells was confirmed by the detection of Dcp1 in exactly these cells, which is a hallmark of apoptosis (Fig. 3E-F). Apoptosis was induced to an even greater extent (and faster) in progenitor cells of the larval tracheal system, where Dcp1-positive cells were observed on day 1 after induction of *Dome.DN* expression (Fig. 3G-I).

**Figure 3:**
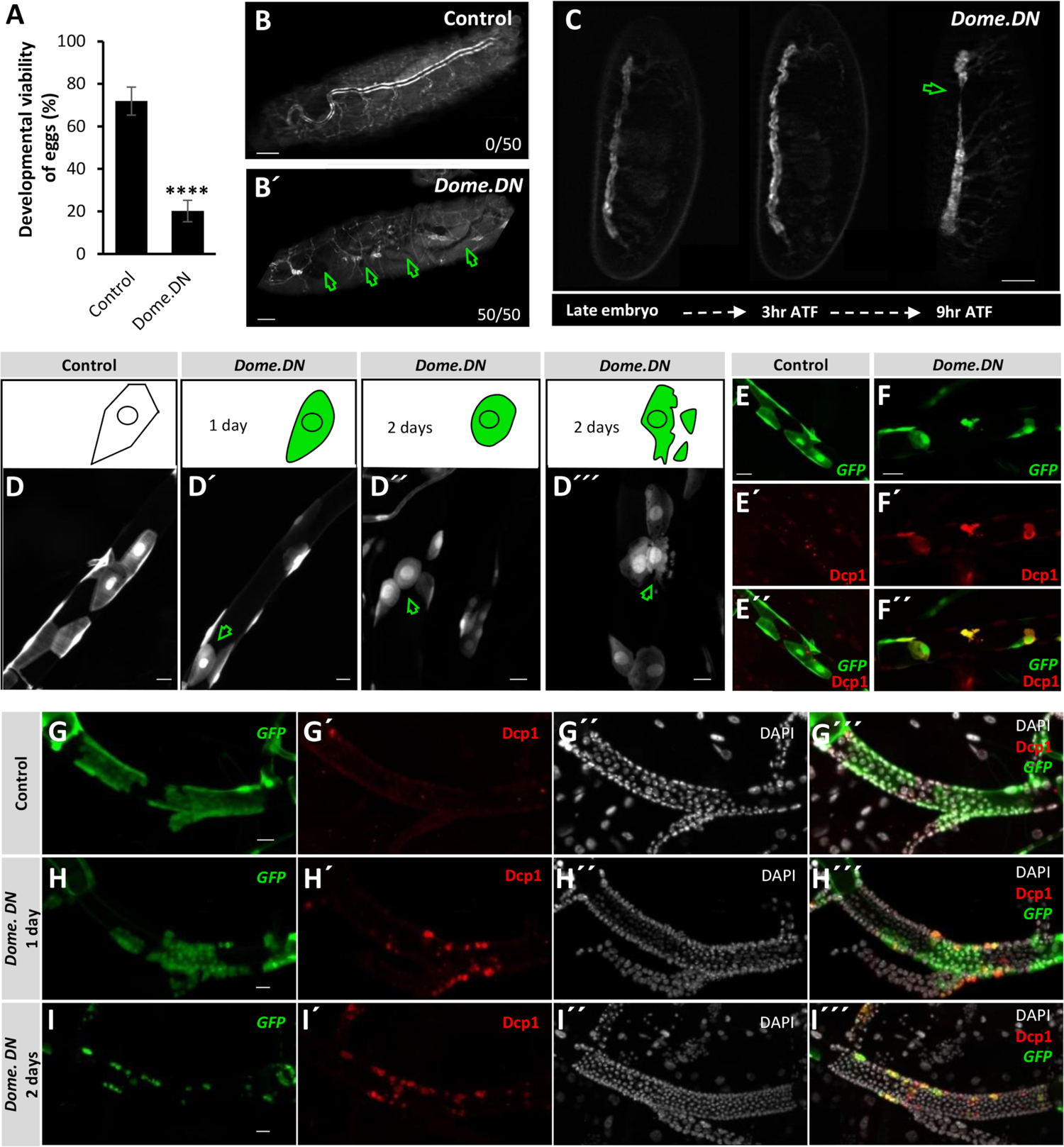
Inhibition of JAK/STAT signaling pathway induced epithelial cell apoptosis. (A) Ectopic blocking of JAK/STAT signaling in the trachea was done by expressing *Dome.DN* under the control of *btl-Gal4*. The developmental viability is shown by the percentage of embryos that hatched. (**** *p* < 0.0001 by Student’s t-test). (B) Micrographs of surviving larvae. Scale bar: 50 µm. *Dome.DN* expression inhibited the formation of an intact tracheal tree. X/XX: emerged phenotype/total animals. (C) Time-lapse imaging of the trachea of *btl-Gal4>Dome.DN* embryos continuously for 9 hours after tracheal formation (9hr ATF). Scale bar: 50 µm. (D-I) Fluorescence micrographs of *Dome.DN* expressing mosaic mutant clones (GFP, green) in the larval trachea (*vvl-FLP*, *CoinFLP-Gal4, UAS-EGFP; tub-Gal80[ts]* (*vvl-coin.ts*)*>Dome.DN*). (D) Negative regulation of JAK/STAT pathway by expressing *Dome.DN* in the trachea led to changes in cell morphology (D’ and D’’) and disintegration afterwards (D’’’, green arrows, Scale bar: 20 µm). (E-F) Fluorescence micrographs of the tracheal somatic cells of *vvl-coin.ts* larvae (E), and the tracheal somatic cells of *vvl-coin.ts>Dome.DN* larvae (F) stained for cleaved Dcp1 (red, on day 2 after induction). Scale bar: 50 µm. (G-I) The progenitor cells of Tr2 could be stained by cleaved Dcp1 earlier than the mutant somatic cells (red, on day 1 after induction). Scale bar: 50 µm. Nuclei are stained with DAPI.

### Increased JAK/STAT signaling induced cell-autonomous structural changes in epithelial cells

We tested the effects of ectopic activation of the JAK/STAT pathway induced by expression of the ligand (*upd3*) or a constitutively active JAK-allele (*Hop.CA*) on tracheal development. These animals died prematurely at the embryo or larval stage (Fig. 4A-B). Microscopic analysis of larvae and embryos showed that the tracheal segment separated at the larval stage (Fig. 4E), however, the disconnection of segments was due to the failure of their fusion at the beginning of tracheal development (Fig. 4C and 4D). In contrast with animals that ectopically expressed *upd3*, more animals expressing *Hop.CA* exhibited an intact tracheal tree (Fig. 4D). However, they all died at the first larval stage, which indicates that JAK/STAT signaling can influence both tracheal formation and growth.

**Figure 4:**
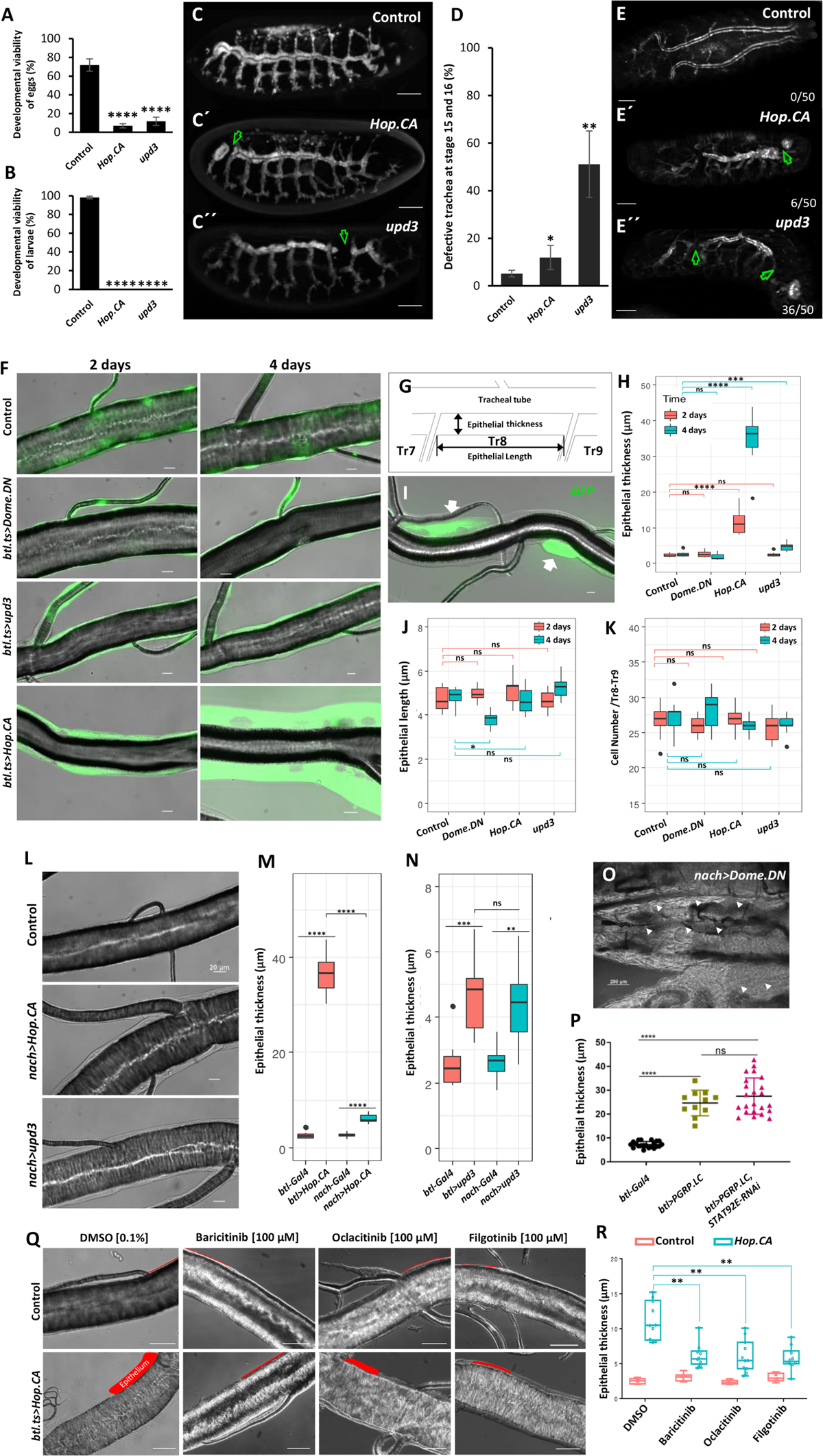
Ectopic activation of JAK/STAT signaling in the trachea induced cell thickening. Ectopic regulation of JAK/STAT signaling in the trachea was performed by expressing *Hop.CA* or *upd3* for activation and *Dome.DN* for inhibition of the pathway, under the control of *btl-Gal4*. The percentage of embryos that hatched (A) and the percentage of larvae that developed into pupa (B) are shown. (C) Fluorescence micrographs of *btl>Hop.CA* (or *btl*>*upd3*) embryos. Scale bar: 50 µm. (D-E) Activation of the JAK/STAT pathway in the trachea led to uncompleted tracheal development during tracheal morphogenesis. (D) Quantification of the numbers of defect trachea in *btl>Hop.CA* or *btl*>*upd3* embryos. (E) Micrographs of the trachea of the surviving larvae. Scale bar: 50 µm. (F) Ectopic activation of epithelial JAK/STAT signaling (*Hop.CA* or *upd3*) or inactivation (*Dome.DN*) under control of *btl.ts-Gal4* (dorsal trunk of Tr8 of L3 larvae). Larvae were raised at 29 ℃ for 2 or 4 days for induction, respectively. Scale bar: 20 µm. (G) Illustration of epithelial thickness, epithelial length. (H) Statistical analysis of the epithelial thicknesses in DT8 of the corresponding larvae. (I) Micrographs of DT8 of *vvl-FLP*, *CoinFLP-Gal4, UAS-EGFP* (*vvl-coin*)*>Hop.CA* larvae. Scale bar: 20 µm. The thickening of the tracheal epithelium is observed in those cells that express *Hop.CA* (arrows). (J-K) Quantification of epithelial length and the cell number of the DT8 region of larvae with different types of ectopic manipulation (including *Dome.DN*, *Hop.CA* and *upd3* expression in the trachea driven by *btl-Gal4, tubPGal80ts*) for 2 or 4 days, respectively. Mild activation of epithelial JAK/STAT signaling mitigated the thickening phenotype. (L) Micrographs of the DT8 regions of *nach-Gal4* larvae, *nach*>*Hop.CA* larvae and *nach>upd3* larvae. Scale bar: 20 µm. (M-N) Quantification of the epithelial thickness of the DT8 region of those larvae whose JAK/STAT signaling were activated by expressing *Hop.CA* or *upd3* under the control of *nach-Gal4* and *btl-Gal4*. (O) Micrographs of *nach>Dome.DN* larvae experiencing suppression of the JAK/STAT pathway make trachea translucent and filled with fluids. Triangle indicates the tracheal position. Scale bar: 200 µm. (P) Quantification of the epithelial thickness of the DT8 region of *btl-Gal4* larvae, *btl>PGRP.LC* larvae and *btl>PGRP.LC, STAT92E-RNAi* larvae. *TubPGal80ts* was used to inhibit the expression of *UAS-PGRP.LC* and *UAS-STAT92E^RNAi^* before animal become L3 larvae and activated *UAS-* genes expression for one day. Changes in epithelial thickness could be rescued by application of specific JAK inhibitors (Q and R). (Q) Microscopy of the tracheal epithelium (L3 larvae). Control crossings *btl-Gal4, tubPGal80ts>w^1118^* compared to JAK/STAT activated crossings *btl-Gal4, tubPGal80ts>Hop.CA.* Scale bar: 50 µm. (R) Quantification of epithelial thickness (highlighted in red) of the JAK inhibitors (Baricitinib, Oclacitinib, Filgotinib) compared to DMSO control. Green arrow: the defective region, ns means no significant, * *p* < 0.05, ** *p* < 0.01, *** *p* < 0.001, **** *p* < 0.0001 by Student’s t-test.

To elucidate the effects of JAK/STAT upregulation in the tracheal system, we employed the temperature inducible Gal4/Gal80[ts] system which could initiate activation of the signaling pathway at different time points during larval development (Fig. S3A). We first tested the effects of inhibition of the tracheal JAK/STAT pathway induced by ectopic expression of *Dome.DN* on the animals. Ectopic expression was initiated at three time points during larval development (Fig. S3B). Larval mortality was higher, the earlier expression was initiated. No larvae survived to the pupal stage (Fig. S3C). On the other hand, activation of JAK/STAT signaling induced by ectopic expression of *upd3* or *Hop.CA* led to lower levels of larval death (Fig. S3B). In both cases, no pupae developed into adults (Fig. S3C). Furthermore, we analyzed the effects of this intervention on tracheal structure. Whereas *Dome.DN* overexpression elicited only minor effects on tracheal structure, *upd3* or *Hop.CA* overexpression caused substantial epithelial thickening (Fig. 4F). Quantitative measurements revealed that ectopic overexpression of *Hop.CA* greatly increased epithelial thickness up to more than 10-fold (Fig. 4H). Experiments involving mosaic expression of *Hop.CA* driven by *vvl-FLP*, *CoinFLP-Gal4, UAS-EGFP (vvl-coin)* demonstrated that this effect was cell-autonomous. In the same trachea, cells that expressed *Hop.CA* were much thicker than their neighboring cells that did not express *Hop.CA* (Fig. 4I). The thicker epithelium could be caused by increased cell volume, increased cell number, or reduced tube surface area. To elucidate the mechanisms underlying this structural response, we analyzed the phenotype in more detail. Consequently, the length-thickness (Fig. 4G) product mirrored the findings made when assessing thickness (Fig. 4J). However, the number of cells in the dorsal trunk of Tr8, or DT8 of the trachea with manipulated JAK/STAT signaling was like that in matching controls (Fig. 4K). Hence, the thickening was due to an increase in cell volume, not an increase in cell number. To determine if this thickening is an artifact caused by the isolation method, we directly fixed the trachea *in situ* or isolated it in cell culture medium with the same osmolarity as the hemolymph. The same thickening was observed in both cases (Fig. S4). To quantify the time course of this thickening, we subjected the trachea to different treatment regimens prior to analysis. At least 2 days of ectopic expression were necessary to induce this phenotype (Fig. S5).

The effects of activating the JAK/STAT pathway on the epithelium were time-dependent (Fig. 4F). To determine if a weaker expression of *Hop.CA* induces similar phenotypes, we used another trachea-specific Gal4-driver, namely, *nach-Gal4* (also called *ppk4-Gal4*). In the *nach-Gal4* line, *Gal4* is expressed from the late embryo stage. Moreover, it is strongly expressed in the early L1 and late L2 stages, at which point expression is stronger than that driven by *btl-Gal4* but is weakly expressed at other larval stages (Liu et al., 2003; Wagner et al., 2009). Microscopic analysis of the trachea showed that weak expression of *Hop.CA* in the trachea promoted epithelial thickening (Fig. 4L). However, epithelial thickening after 4 days of *Hop.CA* induction was lower in *nach>Hop.CA* than in *btl>Hop.CA* (Fig. 4M). On the other hand, weakly driven *upd3* expression in the trachea resulted in identical increases in the thickness as those observed using *btl-Gal4* (Fig. 4N). Finally, we used the same driver to inhibit JAK/STAT signaling in the trachea by expressing *Dome.DN*. Animals died prematurely at the larval stage and their trachea were filled with liquid, demonstrating that functionality was strongly impaired (Fig. 4O). As activation of the IMD pathway by PGRP-LC overexpression induced a very similar phenotype (Wagner et al., 2021), we tested the hypothesis that both, IMD- and JAK/STAT-activation act in the same pathway. Therefore, we activated the IMD-pathway (via PGRP-LC overexpression) while concurrently silencing JAK/STAT signaling (via STAT92e-RNAi) and found no rescue of the thickening phenotype (Fig. 4P).

Treatment with various JAK inhibitors (Roskoski, 2016) reversed epithelial thickening induced by ectopic overexpression of *Hop.CA* in the trachea (Fig. 4Q). Treatment with Baricitinib, Oclacitinib, or Filgotinib reduced epithelial thickening by more than 60% therewith showing its potential in interfering with the JAK/STAT signaling pathway and the induced pathological situations (Fig. 4R).

### Transcriptomic response to enhanced JAK/STAT signaling

To investigate the molecular mechanisms underlying the effects of chronic JAK/STAT activation on the airways, we performed mRNA-sequencing analysis of tracheal cells expressing *Hop.CA* in third instar larvae. The validity of this experimental procedure was controlled by measuring transcript levels of of *hopscotch* (including *Hop.CA*) and *Socs36E* (one of the best-characterized STAT92E target genes) (Bach et al., 2007), which were upregulated 35-fold and 2.7-fold, respectively (*p* < 10^-12^) (Fig. S6B). Principal component analysis of biological replicates separated control and overexpression samples into different groups (Fig. S6A, inner region). Finally, the analysis revealed that the expression of 2004 genes was statistically significantly regulated (*p* < 0.05; > 1.5 fold up or down), with 1128 downregulated genes and 876 upregulated genes (*p* < 0.05). A circular heatmap of a subgroup of them under more stringent conditions with 707 differentially expressed genes (*p* < 0.01 and fold change > 2) is shown (Fig. S6A). Moreover, we analyzed changes in expression upon ectopic silencing of the JAK/STAT pathway using *Dome.DN*. A Venn diagram revealed that there was a statistically significant overlap in the cohorts of genes regulated by both interventions (Fisher’s exact test, *p* < 0.0001; Fig. S6C).

To further elucidate the molecular mechanisms functioning in the thickened epithelium, we performed promotor scan studies and functional enrichment analyses. Genes with the highest rates of induction supported by the lowest p-values were subjected to promoter scanning (Pscan) analysis. Putative genes directly regulated by the JAK/STAT pathway were identified (Tab. 1 and Tab. 2). Among these highly regulated genes, the largest group contained genes that encoded products involved in innate immunity, including seven antimicrobial peptide genes (*IM1 (BomS1), IM2 (BomS2), IM4 (Dso1), IM14 (Dso2), CG5791 (BomBc3), CG5778 (BomT3),* and *Drs*). All these antimicrobial peptides, except for Drs, contain one or two CXXC regions. These CXXC-containing peptides belong to the family of Bomanins, whose expression is highly induced following bacterial or fungal infection, and high levels of the corresponding mature peptides are found in the hemolymph of infected flies (Lindsay et al., 2018).

**Table 1.** Genes that were highly upregulated in response to *Hop.CA* overexpression.

| Name | Maximum group mean | Fold change | FDR p-value |
| --- | --- | --- | --- |
| CG16772 | 11.85 | 135.44 | 0 |
| hop | 286.06 | 35.43 | 0 |
| CG14570 | 42.21 | 35.37 | 0 |
| CG14147 | 55.31 | 32.88 | 0 |
| Hsp70Bb | 161.51 | 29.99 | 0 |
| Ubx | 200.46 | 23.51 | 0 |
| pre-mod(mdg4)-I | 1.12 | 18.67 | 2.22648E-06 |
| IM14 | 11.46 | 15.14 | 3.80132E-07 |
| IM4 | 26.43 | 14.97 | 1.74645E-12 |
| CG14569 | 1576.88 | 14.73 | 0 |
| IM2 | 7.26 | 14.47 | 2.01792E-06 |
| CG5791 | 1.79 | 13.80 | 2.90502E-05 |
| CG9121 | 3.41 | 11.54 | 0 |
| IM1 | 7.16 | 10.97 | 7.52122E-05 |
| CG11413 | 35.70 | 10.64 | 3.44242E-07 |
| CG16710 | 2.13 | 10.32 | 1.72352E-07 |
| CG31960 | 267.26 | 10.05 | 0 |
| CG16789 | 1.45 | 9.46 | 5.58576E-09 |
| CG6034 | 6.92 | 8.47 | 4.7943E-11 |
| Ace | 2.06 | 8.19 | 0 |
| CG13059 | 9.90 | 7.58 | 1.33893E-06 |
| CG31809 | 8.64 | 7.39 | 0 |
| CG7203 | 9.41 | 7.21 | 8.66989E-09 |
| Vha14-2 | 2.51 | 7.05 | 2.31024E-07 |
| wb | 7.35 | 6.69 | 0 |
| Drs | 8166.47 | 6.64 | 0 |
| CG5778 | 3.20 | 6.61 | 0.000294835 |
| CG3457 | 2.69 | 6.34 | 2.38755E-06 |
| Lcp65Ad | 9.93 | 5.79 | 2.58249E-08 |
| lectin-22C | 29.06 | 5.67 | 6.88538E-06 |
| fuss | 3.97 | 5.67 | 0 |
| CG43333 | 2.39 | 5.63 | 1.48392E-09 |
| CG18336 | 2.96 | 5.49 | 5.52167E-05 |
| CG32563 | 3.06 | 5.35 | 0.000696584 |
| FASN2 | 1.65 | 5.34 | 2.95545E-10 |
| Cyp4d21 | 45.70 | 5.31 | 0 |
| Obp57d | 1.80 | 5.28 | 0.000102926 |
| CG17560 | 2.31 | 5.22 | 0.000766701 |
| CG5888 | 68.16 | 5.20 | 0 |

$p < 1 \times 10^{-4}$ , fold change > 5, maximum group mean > 1. The list is restricted to 39 genes.

**Table 2.** Transcription factor-binding site motifs enriched in 41 highly upregulated genes.

| Matrix ID | Matrix name | P-value |
| --- | --- | --- |
| MA0023.1 | dl (var.2) | 0.000443261 |
| MA0022.1 | dl | 0.00407113 |
| MA0532.1 | STAT92E | 0.00794732 |
| MA0450.1 | hkb | 0.00811969 |
| MA0197.2 | nub | 0.0154201 |
| MA0242.1 | Bgb::run | 0.0371696 |
| MA0204.1 | Six4 | 0.0443521 |
| MA0444.1 | CG34031 | 0.0488169 |

A total of 133 transcription factor profiles were used.

Next, we analyzed all significantly regulated genes based on KEGG and GO annotations. Six KEGG pathways and ten categories in GO analysis were enriched in the comparison of the *Hop.CA* expressing trachea compared with the matching controls (Tab. S1 and Fig. 5A-B). One striking feature of the epithelium with chronically activated JAK/STAT signaling was the induced expression of genes involved in “vesicle-mediated transport processes” (Fig. 5A). GO enrichment analysis demonstrated that this term was shared by 32 regulated genes. All these genes were upregulated, except for one (*CG5946*). However, according to a Pscan analysis, these genes did not seem to be direct JAK/STAT targets since predicted STAT92E promoter binding sites were not enriched in the corresponding promoter regions (Tab. S2 and S3). The transcript levels of genes that could be assigned for the term “epicuticle development” were reduced. Here, fifty-five genes relevant to cuticle development were regulated, of which 40 were downregulated. In addition, other biological processes such as “muscle cell differentiation”, “cellular protein-containing complex assembly”, “RNA processing, translation”, and “detection of chemical stimuli” were significantly enriched. In the latter three categories, the number of upregulated genes was like the number of downregulated genes. A schematic representation of the different biological processes superimposed on a scheme of an epithelial cell is shown (Fig. 5B). These alterations at the transcript level were reminiscent of the finding in allergic airways disease, where the expression of cell adhesion molecules or their transport to the membrane of airway epithelial cells are impaired (Bonnelykke et al., 2014; Giridhar et al., 2016; Heijink et al., 2020; Xiao et al., 2011). To test this, we analyzed mouse lungs of an experimental asthma model based on the usage of OVA for sensitization (Fig. 5C-E). Those animals of the asthma model displayed typical structural changes in the airways (Fig. 5D) and the relevant proinflammatory cytokines showed the anticipated increase in expression (Fig. 5E). Consistent with the effects observed in *Drosophila*, the expression of genes involved in the biological process vesicle transport were significantly affected in those mice with an induced experimental allergic airway disease (Fig. 5F) and most of these differentially expressed genes were increased.

**Figure 5:**
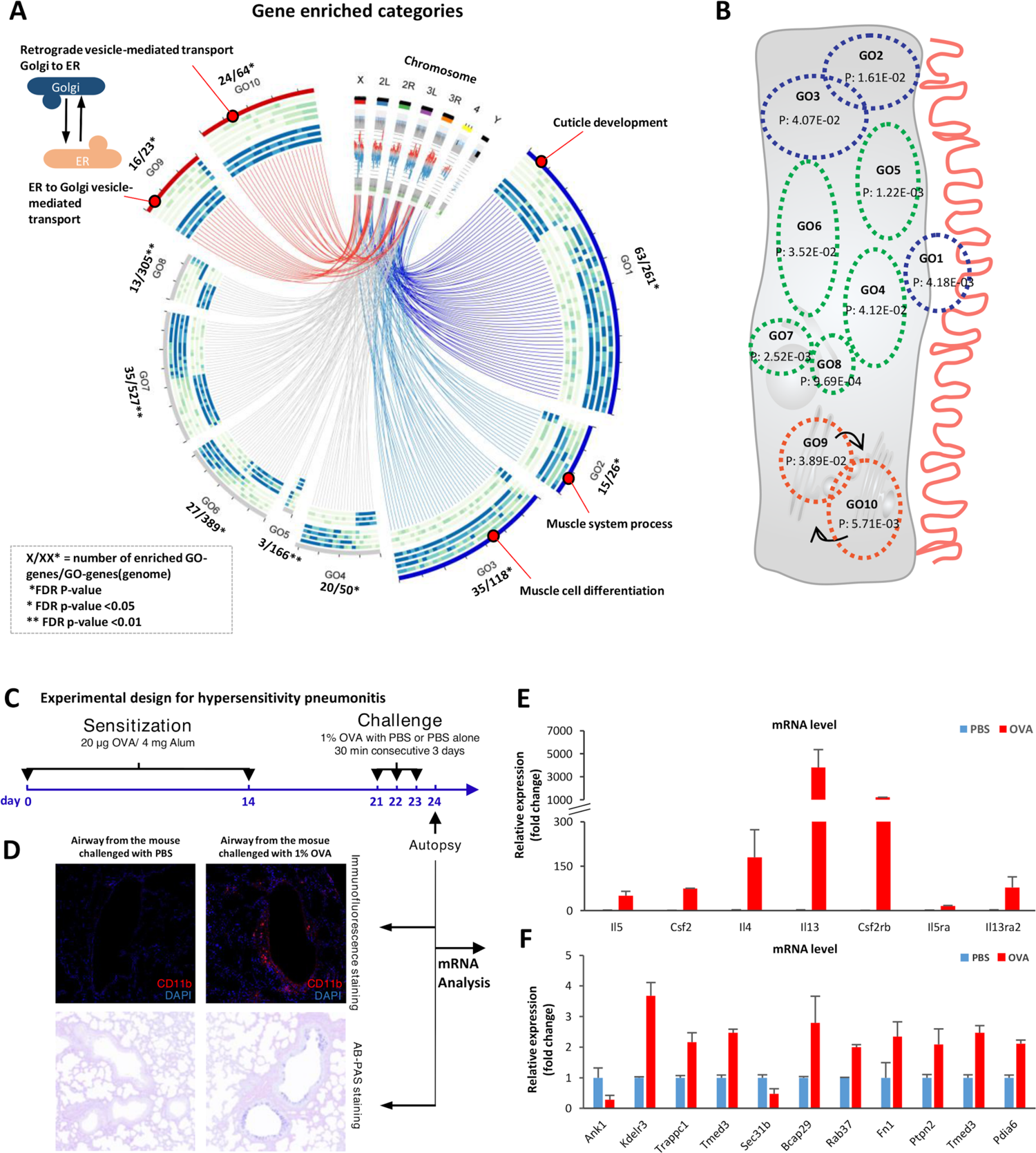
Gene Ontology (GO) enrichment analysis of the 2004 annotated differentially expressed genes in *Hop.CA* overexpressing airways *vs*. controls. (A) Gene ontology analysis showed 10 biological processes that were enriched in *Hop.CA* overexpressing airways compared with matching controls. Most regulated genes in the GO1, GO2, and GO3 were downregulated. Most regulated genes that involved in the COPI-and COPII-mediated vesicular transport between endoplasmic reticulum and Golgi (GO9 and GO10) were up-regulated. GO1, cuticle development; GO2, muscle system process; GO3, muscle cell differentiation; GO4, glutathione metabolic process; GO5, detection of chemical stimulus; GO6, cellular protein-containing complex assembly; GO7, RNA processing; GO8, translation; GO9, endoplasmic reticulum to Golgi vesicle-mediated transport; GO10, retrograde vesicle-mediated transport Golgi to the endoplasmic reticulum. (B) Enriched biological processes were superimposed on a sketch depicting a tracheal epithelial cell, with the corresponding *p*-value added. Blue indicates processes where genes were mainly down-regulated; red indicates processes where genes were mainly up-regulated; green indicates processes where genes were both down-regulated and up-regulated to similar extents. (C) Experimental design for the OVA induced asthma model. Mice were sensitized with intraperitoneal injections of 20 µg of OVA emulsified in aluminum hydroxide in a total volume of 1 ml on days 7 and 14, followed by 3 consecutive challenges each day by exposure to OVA or PBS aerosol for 30 min. Mice were sacrificed 24 h following the final challenge. The left lungs were collected for histological analysis and the superior lobes were dissected for RNA analysis. (D) The lungs were stained with CD11b antibody and with the standard Alcian blue (AB) method followed by the Periodic acid– Schiff (PAS) technique. (E) Important chemokines and chemokine receptors induced by OVA aerosol are involved in JAK/STAT signaling pathway. (F) The genes that function in the process of ER to Golgi vesicle-mediated transport are mostly upregulated when the mice were exposed to OVA aerosol.

### Impairment of protein transport in airway epithelial cells with chronically enhanced JAK/STAT signaling

To evaluate if the transport capacities in the *Drosophila* airway epithelial cells is improved or impaired, we focused on two peripheral membrane-associated proteins involved in cell-cell interactions. These proteins were Coracle (Cora), a component of the septate junctions, and Armadillo (Arm), a component of the adherens junctions. Both proteins depend on vesicle transport to reach their final destinations (Lock and Stow, 2005; Oshima and Fehon, 2011) (Fig. 6A). Immunofluorescence analysis of *Hop.CA-*overexpressing animals revealed that Arm and Cora accumulated in the cytoplasm, which is indicative of dysfunctional vesicle transport (Fig. 6B–E).

**Figure 6:**
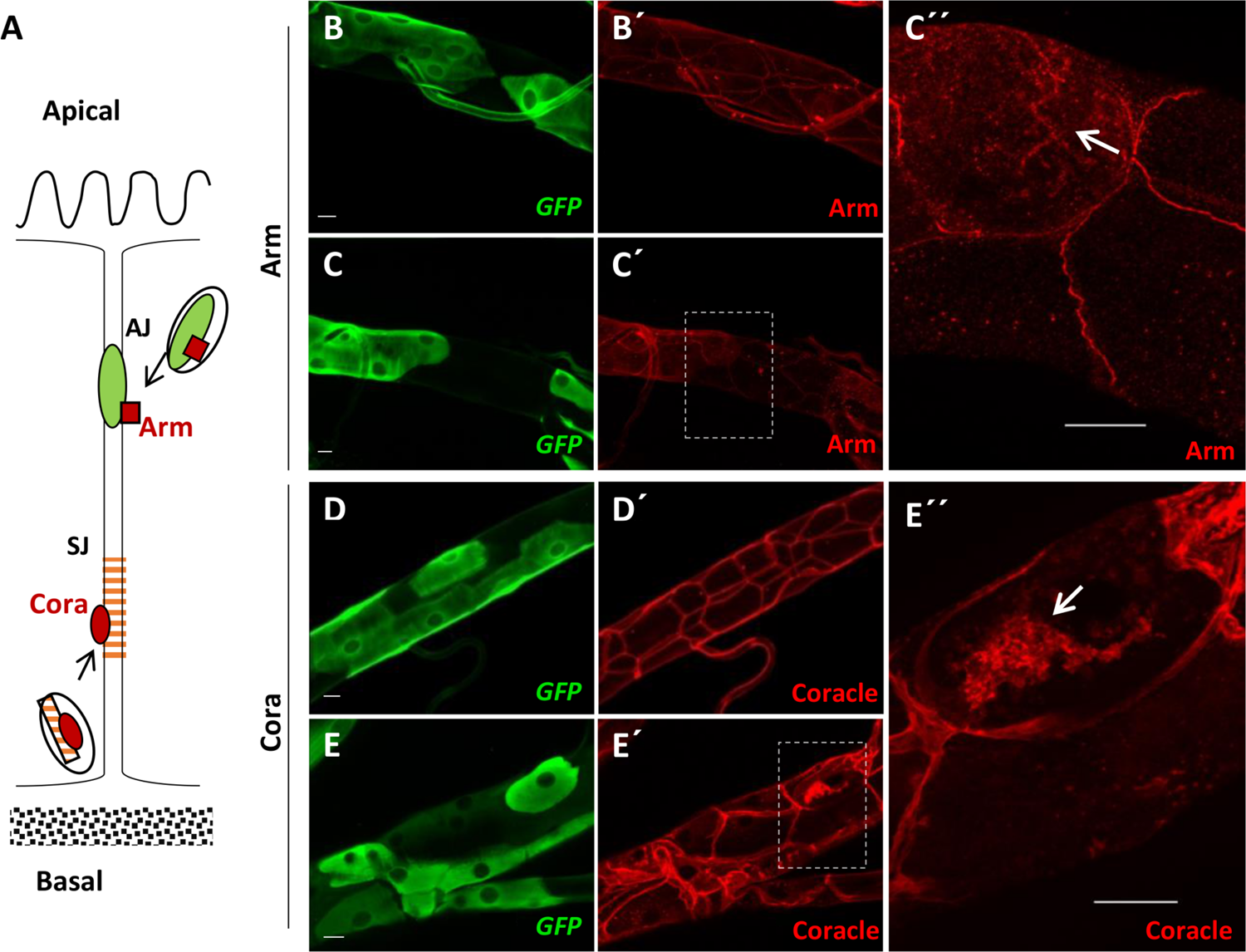
Activation of airway epithelial JAK/STAT signaling affects vesicle-mediated transport of proteins. (A) Schematic orientation of Arm and Cora at the contact zone between two airway epithelial cells. Adherens junction (AJ), septate junction (SJ). Trachea of *vvl-coin* larvae (control; B and D) and *vvl-coin>Hop.CA* larvae (treatment; C and E) stained for GFP (green, Gal4 positive cells), Arm or Cora (red), respectively. 30 specimens were investigated in each group. Scale bar: 20 µm.

The transcriptome data revealed that “cuticle development” was another GO that was enriched. This process depends on the secretory abilities of epithelial cells in the larval trachea. Here, chitin-rich structures, called taenidia, are believed to make tubes flexible as well as sufficiently strong in order to avoid collapse (Glasheen et al., 2010). The chitin structure in the trachea showed a strongly reduced degree of regularity in *Hop.CA*-expressing animals compared with control ones (Fig. 7A-B). Chitin staining revealed that the highly organized structure of the chitinous intima was almost completely lost in *Hop.CA*-expressing animals (Fig. 7A’’ and 7B’’). The mosaic analysis demonstrated that the trachea was structurally disorganized exactly at those sites where *Hop.CA* was expressed (Fig. S7).

**Figure 7:**
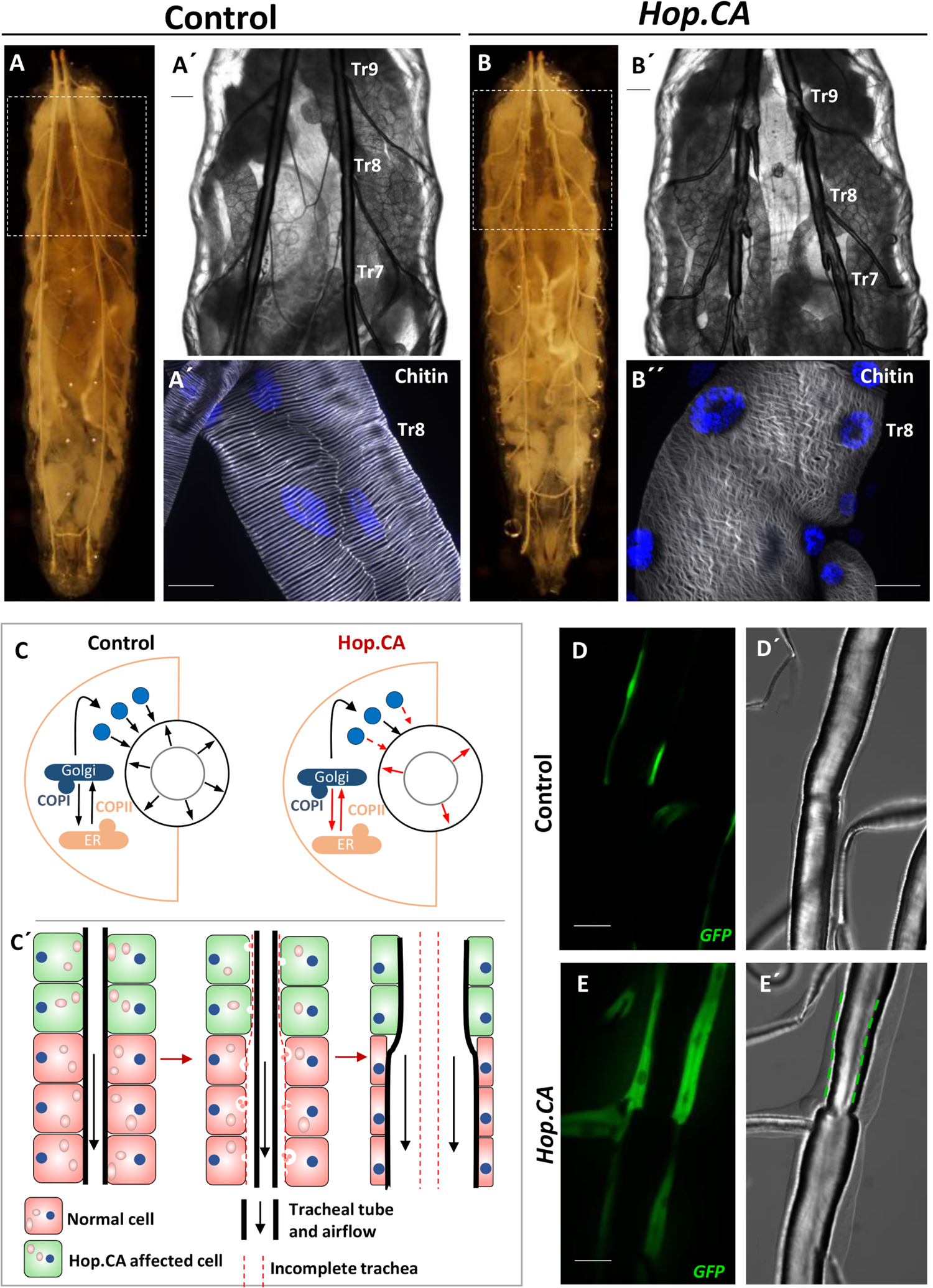
Activation of JAK/STAT signaling in airway epithelial cells affects the development of the epicuticle. (A-B) Activation of JAK/STAT signaling in airway epithelia observed in whole larvae by expressing *UAS-Hop.CA* under the control of the *btl.ts* (B-B’’) compared to the control (A-A’’). In (A’ and B’) the region containing Tr7-Tr9 is shown. (A’’ and B’’) Chitin staining of isolated airways (at the Tr8 region). Scale bar: 50 µm. (C) Schematic illustration of a potential explanation for the thickening of the epithelium and narrowing of the tube. In the control epithelium, a secretory burst of luminal proteins drives the diametric expansion of the tubes, and this process depends on vesicle-mediated transport. COPI-and COPII-mediated vesicular transport plays a central role in this process, mutation members in the process underly the tube size defect in previous reports (Jayaram et al., 2008; Tsarouhas et al., 2007). Although active JAK induced the expression of the genes involved in the vesicle-mediated transport, long-term activation eventually impedes the transport of vesicles out of the cell (C). This dysfunction of transport could lead to the appearance of tube size defects such as the increase in the cell volume and narrowing of the tube size (C’) (Jayaram et al., 2008; Tsarouhas et al., 2007). (D-E) Micrographs of the trachea of *vvl-coin* larvae (control; D) and *vvl-coin>Hop.CA* larvae (treatment; E). Cells with *vvl-Gal* expression are stained with GFP (D-E). The green dash line in the E’ show stenosis in the regions with cells expressing *Hop.CA* to a high degree. Scale bar: 50 µm.

We propose a simple model that explains structural phenotypes mainly based on the impaired vesicle-mediated transport and the lack of adhesion-associated proteins to the membrane, which is in line with previous findings about this biological process (Jayaram et al., 2008; Tsarouhas et al., 2007) (Fig. 7C). Lumen narrowing is another phenotype that might be caused by the dysfunction of the process which could be observed in our study as well. This is prominently seen when epithelial cells expressed *Hop.CA* at very high levels, as indicated by strong green fluorescence (Fig. 7D-E and Fig. S4).

## Discussion

The present study aimed to clarify the generic role of the JAK/STAT pathway in the respiratory epithelium. We mainly used the fruit fly *Drosophila* because it has ideal characteristics for conducting this study. The unique features of this model allowed us to focus on the essentials, namely the entirety of the signaling pathway and its importance in respiratory epithelial cells. The JAK/STAT signaling pathway in mammals has a degree of redundancy and parallelism in all its levels, rarely observed in other signaling systems. However, this exceptionally high degree of parallelism hampers determing the significance of JAK/STAT signaling per se and whether it differs substantially from signaling through individual cytokines or receptors. As mentioned earlier, the Drosophila JAK/STAT signaling pathway is simple, allowing targeted manipulation. Here, we showed that 1) JAK/STAT signaling exhibited tonic activity in almost all respiratory epithelial cells, 2) that exposure to airborne stressors such as hypoxia or cigarette smoke enhanced its activity, 3) that blocking JAK/STAT signaling induces apoptosis, and that 4) ectopic activation in epithelial cells induced several phenotypes reminiscent of those found in chronic inflammatory lung diseases such as epithelial thickening or narrowing of the airways. The main results 1-3 are thematically very closely related. The tonus of JAK/STAT signaling shown in embryonic, larval, and adult airway epithelia suggests that this cellular function is essential as it probably ensures the survival of these epithelial cells. JAK/STAT signaling is required to maintain the cellular identity of airway epithelial cells (D’Amico et al., 2018) and is important for the response to airway epithelial injury and epithelial cell survival (Juncadella et al., 2013; Paris et al., 2020; Tadokoro *et al*., 2014). This agrees with our studies, which revealed that blockade of the signaling pathway leads to apoptosis of airway epithelial cells of flies. Furthermore, it implies that the JAK/STAT pathway plays a central role in homeostatic processes in this organ, which also appears to be the case in mammalian airway epithelia. Perturbation of JAK/STAT signaling induces apoptosis in cell lines derived from airway epithelia (Zheng et al., 2016), and this is particularly relevant for tissues and cells that can still divide or grow (Quinton and Mizgerd, 2011). This ability to grow might be one reason for the high susceptibility of Drosophila airway epithelial cells to the blockade of JAK/STAT signaling. Thus, cytokines that activate the JAK/STAT signaling pathway act as survival factors, especially when repair mechanisms are operative. Functional JAK/STAT signaling is also required for regenerative processes after epithelial damage (Kida *et al*., 2008; Paris *et al*., 2020; Tadokoro *et al*., 2014), the response to infections (Matsuzaki et al., 2006), and the reaction to hyperoxia (Hokuto et al., 2004). This indicates that JAK/STAT signaling in mammalian airway epithelial cells and the *Drosophila* trachea is particularly relevant for a protective reaction to stressful stimuli. In *Drosophila*, we observed a basal level of JAK/STAT signaling in these cells, and this signaling was further activated in response to very strong stressors such as chronic exposure to smoke particles. An organ-autonomous JAK/STAT signaling system seems to be operative, composed of the cytokines Upd2 and Upd3 and the activated signaling pathway induced in the same regions of the tracheal system. A comparable, stress-induced system was identified in the intestinal epithelium of flies, where highly stressful insults targeting absorptive enterocytes induce the production and release of the cytokine Upd3 (Jiang et al., 2009; Miguel-Aliaga et al., 2018). In contrast to the intestine, where cytokines produced by stressed enterocytes induce the proliferation of stem cells to replenish the enterocyte pool, stem cells are not involved in the tracheal response.

Although a threshold level of JAK/STAT signaling is required for the functionality and survival of airway epithelial cells, excessive activation of this signaling is associated with several lung diseases such as lung cancer, acute lung injury, asthma, pulmonary fibrosis, and COPD (Adnot et al., 2019; Chen et al., 2021; D’Amico *et al*., 2018; Dutta *et al*., 2014; Gao et al., 2004; Milara *et al*., 2018; Parakh et al., 2021; Prele et al., 2012; Simeone-Penney et al., 2007; Yew-Booth *et al*., 2015; Zhang et al., 2012). The ectopic activation of JAK/STAT signaling in the fly’s airway epithelium induced major structural changes, mainly to the architecture of tracheal cells. The mosaic analysis demonstrated that this effect was cell autonomous. These structural changes of the trachea are reminiscent of those observed in asthma, COPD, acute lung injury, and lung cancer. Structural changes that permit fulfillment of the original function of epithelial cells are hallmarks of the epithelial-to-mesenchymal transition. In lung cancer, the epithelial-to-mesenchymal transition depends on JAK/STAT signaling (Liu et al., 2014).

Ectopic JAK/STAT pathway activation also induced complex transcriptomic alterations in airway epithelial cells. These changes also affected immune-relevant genes, such as those encoding antimicrobial peptides, mainly from the Bomanin family. This dependency of antimicrobial responses on JAK/STAT signaling is also seen in the mammalian airway epithelium (Choi et al., 2013; Simeone-Penney *et al*., 2007). However, chronically activated JAK/STAT signaling perturbed secretory processes and the formation of extracellular structures. The extracellular matrix defects caused by *Hop.CA* overexpression are reminiscent of the pathophysiology of the aforementioned chronic lung diseases that involve responses to acute or chronic injury (Fahy and Dickey, 2010). Therefore, attempts to enhance the repair capacities of epithelial cells by inducing structural changes via excessive JAK/STAT signaling would also perturb normal cellular functions, such as the transport of junction proteins to the membrane. This would lead to a reduced barrier function of the epithelium, which is a hallmark of chronic lung diseases such as asthma and COPD (Georas and Rezaee, 2014; Gon and Hashimoto, 2018; Heijink et al., 2012). We also observed epicuticular changes, which considerably influenced the structure of the whole organ. However, it is difficult to identify an equivalent response in vertebrates.

Furthermore, our finding that the JAK/STAT signaling pathway operates in airway epithelial cells of embryos, larvae, and adults implies that it plays a central role in these cells at all developmental stages. The JAK/STAT pathway is involved in the embryonic development of the tracheal system (Hombria and Sotillos, 2013). We clarified the sequence of developmental steps in which this signaling pathway plays a central role. The JAK/STAT signaling pathway is particularly relevant for developing and maintaining the dorsal trunk. It is a matter of debate, whether JAK/STAT signaling is essential for mammalian lung development. However, the prevailing view that JAK/STAT signaling is not essential for embryonic lung development (Tadokoro et al., 2014) has been challenged by recent studies (Piairo et al., 2018). Although JAK/STAT signaling is essential for different aspects of tracheal development, the major focus of the current study was to understand its role in maintaining homeostasis in the fully functional airway epithelium.

The JAK/STAT pathway is an excellent target for therapeutic intervention in many lung diseases, including asthma, COPD, acute lung injury, idiopathic pulmonary fibrosis, and lung cancer (Athari, 2019; Loh et al., 2019; Milara *et al*., 2018; Severgnini et al., 2004; Song et al., 2011; Yew-Booth *et al*., 2015). We demonstrated that pharmacological interference of the JAK/STAT signaling pathway reverted the structural phenotype observed upon ectopic activation of this signaling and, consequently flies survived. This shows that the *Drosophila* model is not only suitable to study the structural effects of excessive JAK/STAT signaling and the underlying molecular mechanisms but also to screen compounds and thereby identify novel therapeutic strategies. Moreover, the intrinsic architecture of the *Drosophila* system allows the prediction that the major site of action is the airway epithelium, thus implying that inhalation would be the ideal route of drug administration in a therapeutic setting.

It should be remembered that the vertebrate lung and insect trachea are not homologous. Nevertheless, both airway systems share a high degree of similarity regarding their development, physiology, innate immunity, and operative signaling systems (Andrew and Ewald, 2010; Bergman et al., 2017). Therefore, the fruit fly is a valuable tool for studying genes associated with a great variety of chronic lung diseases, including asthma, COPD, and lung cancer (Bossen et al., 2019; Kallsen et al., 2015; Levine and Cagan, 2016; Prange *et al*., 2018; Roeder *et al*., 2009; Roeder *et al*., 2012). This simple model can be used as part of an experimental toolbox to elucidate the role of JAK/STAT signaling in the airways and the effects of chronic deregulation of this signaling. In addition, it provides a readily accessible experimental system that is amenable to pharmacologic interventions and allows hypotheses and intervention strategies to be easily tested.

## Material and Methods

### *Drosophila* strains and husbandry

*STAT92E-GFP* was used to monitor the activation of the pathway (Bach et al., 2007); the Gal4-UAS system (Brand and Perrimon, 1993) was used to target ectopic expression to the tracheal system. Gal4 drivers used were: *btl-Gal4, UAS-GFP* on the 3rd chromosome, and *btl-Gal4, UAS-GFP* on the 2nd chromosome (obtained from the Leptin group, Heidelberg, Germany); *upd2-Gal4*; *upd3-Gal4* (Prange et al., 2018); *nach-Gal4* (Liu et al., 2003). The UAS responders included: *UAS-LacZ.nls* (BDSC 3956), *UAS-domeΔcyt2.1* (*UAS-Dome.DN*) and *UAS-Hop.CA* (*UAS-hopTumL*) were obtained from N. Perrimon (Brown et al., 2001; Harrison et al., 1995). The *UAS-upd3* was constructed in our lab. *TubP-Gal80[ts]* (BDSC 7018) was obtained from the Bloomington stock center. Unless otherwise stated, the flies were raised on standard medium at 25 °C with 50–60% relative humidity under a 12:12 h light/dark cycle as described earlier (Fink et al., 2016).

### Coin-FLP expression system

*Vvl-FLP/CyO; btl-moe.mRFP* (BDSC 64233), *tubP-Gal80[ts]* and *CoinFLP-Gal4, UAS-2xEGFP* (BDSC 58751) were used to construct animals for the tracheal mosaic analysis. Ventral veins lacking (*vvl*) was expressed in larval tracheal clones that covered approximately 30 to 80% of the trachea (Bosch *et al*., 2015; Chen and Krasnow, 2014). The genotype of the flies was *vvl-FLP*, *CoinFLP-Gal4*, *UAS-2xEGFP*/*CyO* (*vvl-coin*) and *vvl-FLP*, *CoinFLP-Gal4*, *UAS-2xEGFP*/*CyO*; *tub-Gal80[ts]* (*vvl-coin.ts*).

### Developmental viability

For developmental viability of eggs, the eggs were collected overnight and were not physically handled in any way. The number of the hatched eggs and the pupae were counted starting from two days after the collection. Each group has 4 replicates that included more than 20 eggs. For developmental viability of larvae, *tub-Gal80[ts]* was used to limit UAS responder expression at the larvae stage. Animals were raised at 18 °C to keep the UAS responder gene silent. Larvae at different instar stages were transferred to a new medium at 29 °C. In this study, usually 4 replicates were performed with 30 larvae. The stage of larvae was determined via the appearance of anterior spiracles.

### Determination of epithelial thickness

The trachea of L2 and L3 Larvae were carefully dissected from the posterior side of the body in PBS. The isolated trachea were immersed in 50% glycerol and digital images were captured within 15 min. L2 Larvae were distinguished from L3 larvae by the appearance of anterior spiracles. The relative ages of L3 larvae were inferred from the size of the animal. 30 larvae were used in each group and specimens were analyzed by Image J.

### Drug application in *Drosophila*

JAK inhibitors (Baricitinib #16707, Oclacitinib #18722, Filgotinib #17669 - Cayman Chemicals, Michigan, USA) were diluted in DMSO [100 mM]. For later application the inhibitors were diluted 1:10 in 100% EtOH. We used 20 µl of each diluted inhibitor for 2 ml of concentrated medium (5% yeast extract, 5% corn flour, 5% sucrose, 1% low-melt agarose, 1 ml of 10% propionic acid and 3 ml of 10% Nipagin). The eggs of each crossing have been applicated on the modified medium and kept at 20 °C until the larvae reach the L2 stage. Afterwards they were incubated at 30 °C for 2 days to induce the *btl-Gal4, UAS-GFP; tub-Gal80[ts]* (*btl.ts*)–driver. The trachea of L3 Larvae were carefully dissected from the posterior side of the body in PBS. Isolated trachea was immersed in 50% glycerol and digital images were captured in 15 min. 10 larvae were used for each group.

### Experimental design for hypersensitivity pneumonitis in mice

Experimental design for hypersensitivity pneumonitis contains two processes, sensitization and challenge of mice with ovalbumin (Sigma Aldrich, A5503). The protocol refers to the protocol that (Daubeuf and Frossard, 2013) described with a few modifications. 8 week-old Balb/C mice were sensitized with intraperitoneal injections of 20 µg of OVA emulsified in aluminum hydroxide in a total volume of 1 ml on days 7 and 14, followed by 3 consecutive challenges each day by exposure to OVA or PBS aerosol for 30 min. Mice were sacrificed 24 h following the final challenge. The left lungs were collected for histological analysis and the superior lobes were dissected for RNA analysis. 6 mice per group were used in this experiment and 4 mice were randomly selected in each group for analyses. Total mRNA sequencing, data processing, and statistical analysis were performed by Genesky (Shanghai, China). The experimental protocols were approved by the Animal Care and Protection Committee of Weifang Medical University (2021SDL418).

### AB-PAS staining and immunofluorescence analysis of mouse lung

Mice were sacrificed in excess CO_2_ gas. The lungs of euthanized mice were inflated by intratracheal injection of cold 4% paraformaldehyde and then were fixed and embedded in paraffin as described in (Baligar et al., 2016). AB-PAS staining, and Immunofluorescence analysis were supported by Servicebio (Wuhan, China). All sections were photographed using a microscope slide scanner (Pannoramic MIDI: 3Dhistech). The materials and methods could be found at the following link https://www.servicebio.cn/data-detail?id=3040&code=MYYGSYBG and https://www.servicebio.cn/data-detail?id=3595&code=RSSYBG. We list the main reagents here: AB-PAS solution set (Servicebio, G1049), Anti-CD11b (Servicebio, GB11058) and DAPI (Servicebio, G1012-10ML).

### Time-lapse microscopy

All images were acquired using a ZEISS Axio Image Z1 and a ZEISS LSM 880 fluorescent microscope (INST 257/591-1 FUGG). Embryos were dechorionated in 3% sodium hypochlorite and immersed in Halocarbon oil 700 (Sigma Aldrich, 9002-83-9). Then the embryos were imaged after stage 15 when the tracheal tree formed at 3 hours intervals. 40 embryos were investigated in each group.

### Cigarette smoke and hypoxic exposure

All cigarette smoke exposure experiments were carried out in a smoking chamber, attached to a diaphragm pump. Common research 3R4F cigarettes (CTRP, Kentucky University, Lexington, USA) were used for all experiments. The vials containing animals were capped with a monitoring grid to allow the cigarette smoke to diffuse into the vial. For long-time smoke experiments, L2 larvae were exposed to smoke three times a day for 30 min each, on two consecutive days. For heavy smoke experiments, L3 larvae were exposed to smoke of 2 cigarettes for 45 min, which led to about 35-50% mortality. To study the effects of hypoxia on the activity of JAK/STAT signaling pathway, larvae were exposed to long-term hypoxia and short-term hypoxia separately. For long-term hypoxia experiments, L2 larvae were exposed to 5% oxygen three times a day for 30 min each, on two consecutive days. For short-term hypoxia experiments, L3 larvae were exposed to 1% oxygen once for 5 hours. At least 20 animals in 4 vials were investigated in each group.

### Immunohistochemistry of *Drosophila* trachea

Larvae were dissected by ventral filleting and fixed in 4% paraformaldehyde for 30 min. Embryos were staged according to Campos-Ortega and Hartenstein (Campos-Ortega and Hartenstein, 1997) and fixed in 4% formaldehyde for 30 min. Immunostaining followed standard protocols as described earlier (Jeon et al., 2008; Levi et al., 2006). GFP signals were amplified by immunostaining with polyclonal rabbit anti-GFP (used at 1:500, Sigma-Aldrich, Merck KGaA, Darmstadt, Germany, SAB4301138). 40-1a (used at 1:50, DSHB, Iowa City, USA) was used to detect Beta-galactosidase. Coracle protein was detected with a monoclonal mouse anti-coracle antibody (used at 1:200, DSHB, Iowa City, USA, C566.9). Armadillo protein was detected with a monoclonal mouse anti-armadillo antibody (DSHB, US, N2 7A1, used at 1:500). A monoclonal rabbit Cleaved Drosophila Dcp1 (used at 1:200, Cell Signaling, Frankfurt/M, Germany, #9578) was used to detect apoptotic cells. Secondary antibodies used were: Cy3-conjugated goat-anti-mouse, Cy3-conjugated goat-anti-rabbit, Alexa488-conjugated goat-anti-mouse (used at 1:500, Jackson Immunoresearch, Dianova, Hamburg, Germany), Alexa488-conjugated goat-anti-rabbit (used at 1:500, Cell signaling, Frankfurt/M, Germany, #9578). Tracheal chitin was stained with 505 star conjugated chitin-binding probe (NEB, Frankfurt/M, Germany, used at 1:300). Nuclei were stained with 4’,6-Diamidino-2-Phenylindole, Dihydrochloride (DAPI) (Roth, Karlsruhe, Germany, 6843). 30 specimens were investigated in each group. Specimens were analyzed and digital images were captured either with a confocal (Zeiss LSM 880, Oberkochen, Germany) or a conventional fluorescence microscope (ZEISS Axio Imager Z1, Zeiss, Oberkochen, Germany), respectively.

### RNA isolation and RNA sequencing

For the gene expression analysis of 3rd instar larvae trachea, animals were dissected in cold PBS and isolated trachea transferred to RNA Magic (BioBudget, Krefeld, Germany) and processed essentially as described earlier (Prange et al., 2018) with slight modifications. The tissue was homogenized in a Bead Ruptor 24 (BioLab products, Bebensee, Germany) and the RNA was extracted by using the PureLink RNA Mini Kit (Thermo Fisher, Waltham, MA, USA) for phase separation with the RNA Magic reagent. An additional DNase treatment was performed following the on-column PureLink DNase treatment protocol (Thermo Fisher, Waltham, MA, USA).

Sequencing libraries were constructed using the TruSeq stranded mRNA kit (Illumina, San Diego, USA) and 50 bp single-read sequencing was performed on an Illumina HiSeq 4000 with 16 samples per lane. Resulting sequencing reads were trimmed for low-quality bases and adapters using the fastq Illumina filter (http://cancan.cshl.edu/labmembers/gordon/fastq_illumina_filter/) and cutadapt (version 1.8.1) (Martin, 2011). Transcriptomics analysis including gene expression values and differential expression analysis was done using CLC Genomics Workbench. The detailed protocols can be obtained from the CLC Web site (http://www. clcbio.com/products/clc-genomics-workbench). *Drosophila melanogaster* reference genome (Release 6) (Hoskins et al., 2015) was used for mapping in this research.

The transcription factor binding site enrichment and the Gene Ontology enrichment analyses of the differentially expressed genes were carried out using Pscan (http://159.149.160.88/pscan/) and online GO enrichment analysis (http://geneontology.org/), respectively. We chose-450-50 bases of the annotated transcription start site of the genes as the transcription factor binding sites for enrichment analysis. The analysis performed with the TFBSs matrices that is available in the JASPAR databases (version Jaspar 2018_NR). Data were visualized through the circos software http://circos.ca/software/.

### Statistics and reproducibility

We did not use statistical methods to predetermine sample sizes but the sample sizes used in this study are similar or higher as those used in previous studies (Proske et al., 2021; Wagner *et al*., 2021). Specific approaches to randomly allocate samples to groups were not used and the experiments were not performed in a blinded design. No data were excluded from the analysis. Prism (GraphPad version 7) was used for statistical analyses and the corresponding tests used are listed in the figure legends.

