## Supplementary material for "Maintaining structural and functional homeostasis of the *Drosophila* respiratory epithelia requires stress-modulated JAK/STAT activity"

**Supplementary materials for Niu et al.**

### Supplementary figures

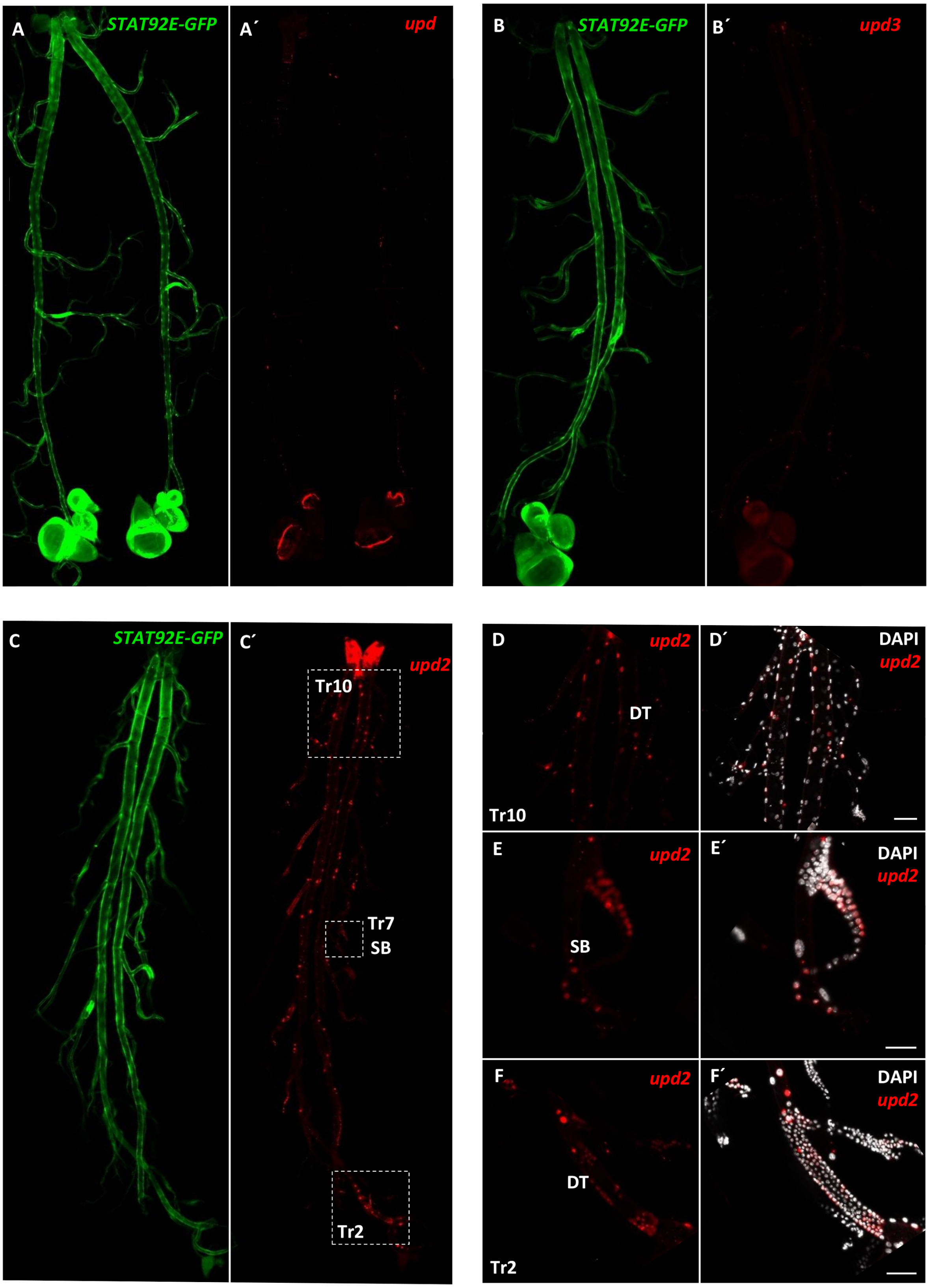

Figure S1 JAK/STAT signaling is activated in the larval trachea. (A-C) Fluorescence micrographs of the trachea of ligands (*upd*, *upd2* or *upd3*)-*Gal4*, *UAS-LacZ.nls*; *STAT92E-GFP* larvae stained to show ligand expressing cells (anti-ß-galactosidase, red) and JAK/STAT signaling activated cells (anti-GFP, green). *Upd2* displayed a higher transcript level in the trachea compared to *upd* (showed a strong expression in the other tissue, like an imaginal disc) and *upd3* (which expression in the trachea could be induced by CS (Fig. 5 and (Prange et al., 2018)). (D-F) Fluorescence micrographs of Tr2 DT, Tr7 SB and Tr10 DT of *upd2-Gal4, UAS-LacZ.nls*; *STAT92E-GFP* larvae. In the dividing active regions such as Tr2 DT and Tr7 SB there is stronger JAK/STAT activity and a high intensity of *upd2* positive cells (E-F) compared to their adjoining somatic cells (D). Nuclei are stained with DAPI. Scale bar: 50 µm.

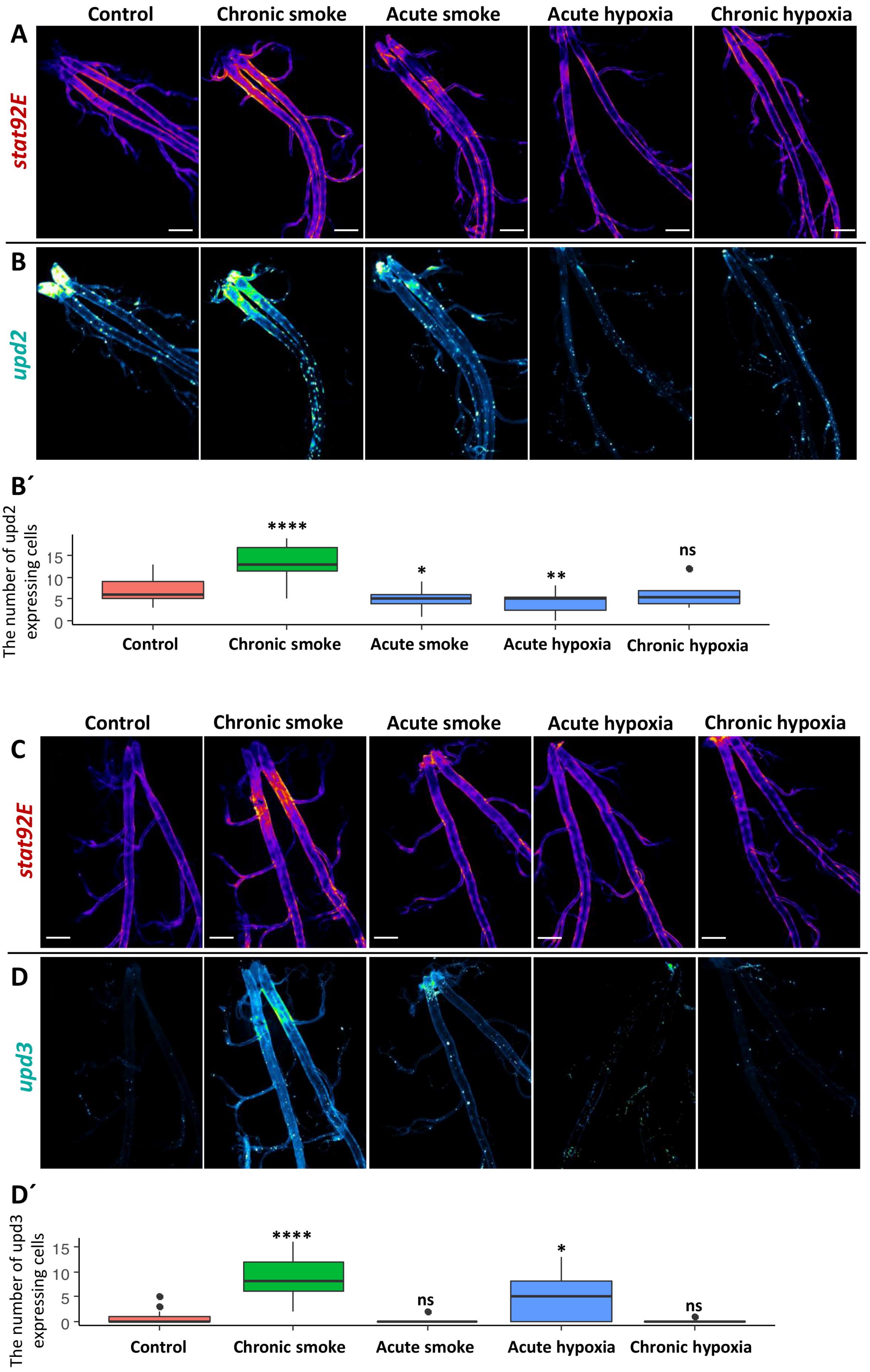
Control
Chronic smoke
Acute smoke
Acute hypoxia
Chronic hypoxia
A
stat92E
B
upd2
B´
The number of upd2 expressing cells
15
10
5
0
****
*
**
ns
Control
Chronic smoke
Acute smoke
Acute hypoxia
Chronic hypoxia
Control
Chronic smoke
Acute smoke
Acute hypoxia
Chronic hypoxia
C
stat92E
D
upd3
D´
The number of upd3 expressing cells
15
10
5
0
****
ns
*
ns
Control
Chronic smoke
Acute smoke
Acute hypoxia
Chronic hypoxia

Figure S2 The activity of JAK/STAT pathway and the expression of its ligands *upd2* or *upd3* were observed in the trachea of larvae that were exposed to smoke and hypoxia. Fluorescence micrographs of the trachea of *upd2-Gal4* (A-B) *or upd3-Gal4* (C-D); *STAT92E-GFP*; *UAS-LacZ.nls* larvae that were exposed for 2 days smoke (chronic smoke), heavy smoke (acute smoke), strong hypoxia (acute hypoxia), and 2 days hypoxia (chronic hypoxia). Trachea were stained for GFP (A and C; red, JAK/STAT pathway activated zones) and Beta-galactosidase (green, cells that expressed *upd2* or *upd3*), respectively. The numbers of cells that expressed *upd2* (B') or *upd3* (D') were counted under these different conditions. ns means not significant, * $p < 0.05$, ** $p < 0.01$, **** $p < 0.0001$ by Student's t-test. Scale bar: 200 µm.

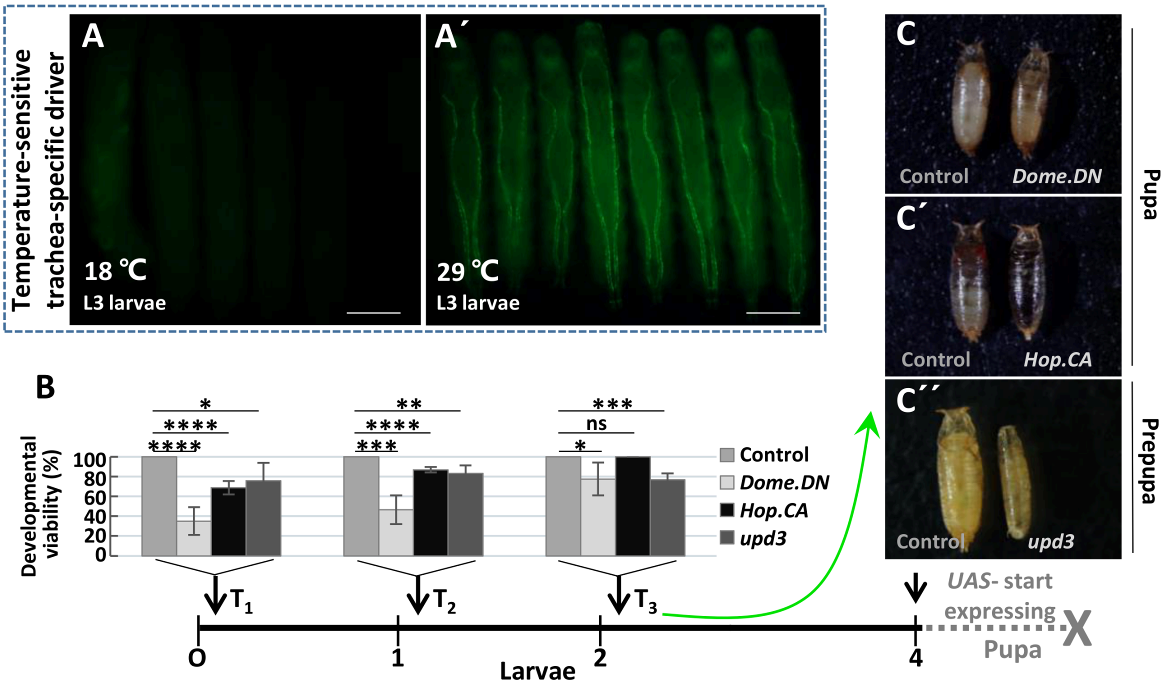


Figure S3 The temperature sensitive tracheal-specific driver line and the developmental viability analysis of larvae with abnormal JAK/STAT signaling. (A) *btl-Gal4, UAS-GFP; tub-Gal80[ts]* (*btl.ts*) larvae were raised at 18 ℃ (nonpermissive, A) and 29 ℃ (permissive, A´), respectively. Using the Gal4/UAS system, comprising the temperature-sensitive repressor Gal80[ts], were used to time ectopic gene expression. (B) Developmental viability of larvae with different genotypes (including *Dome.DN*, *Hop.CA*, and *upd3* expression in the trachea driven by *btl.ts*) and different start points of expression (indicated by black arrows). (C) None of these manipulations allowed survival up to adults shown by pictures of the non-eclosed pupae. ns means not significant, * $p < 0.05$, ** $p < 0.01$, *** $p < 0.001$, **** $p < 0.0001$ by Student's t-test.

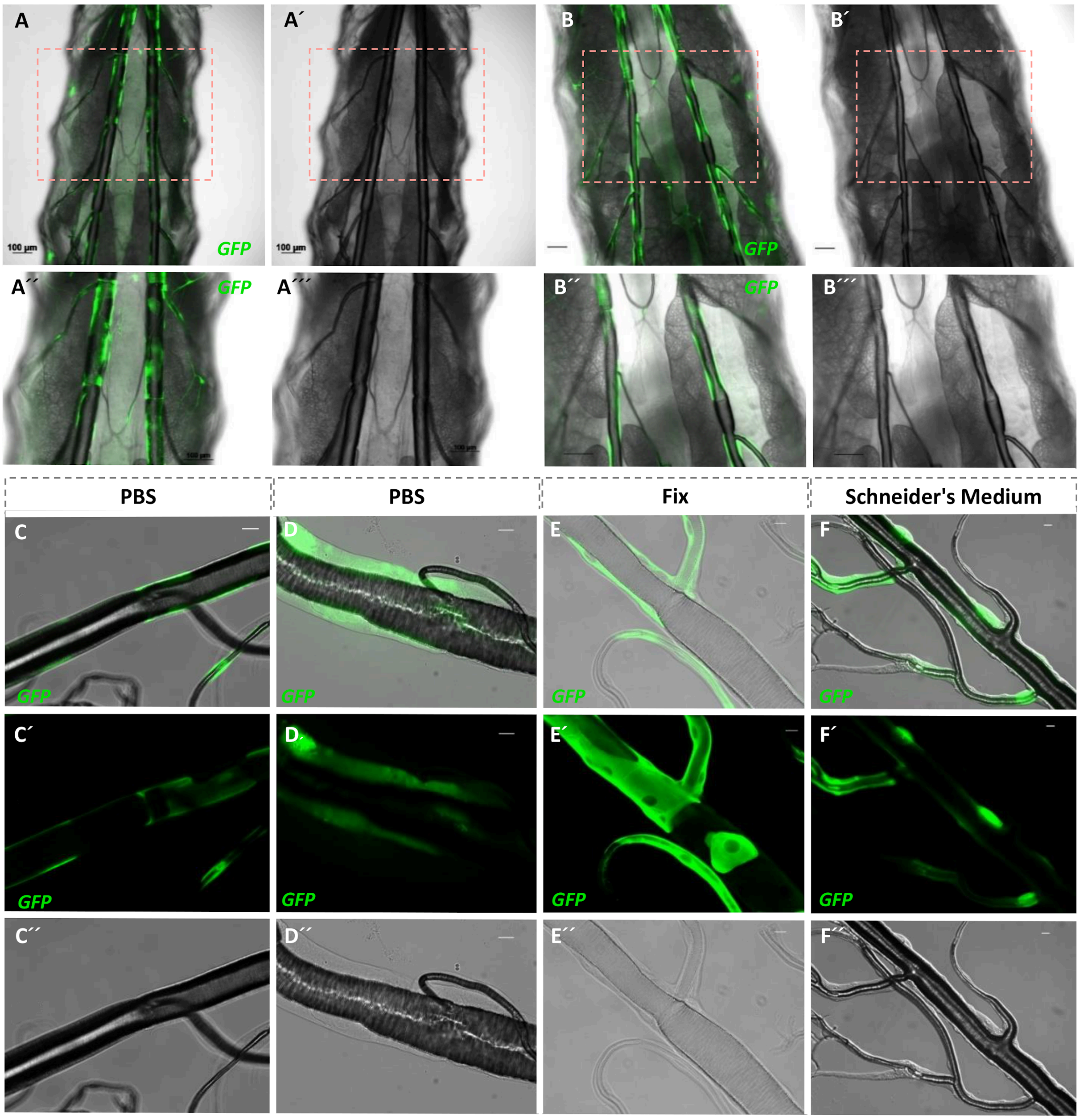


Figure S4: Validation that the increase in cell volume is not an artefact. (A-B) Micrographs of the posterior of *vvl-coin* larvae (A) and *vvl-coin>Hop.CA* larvae (B). Microscopy of the trachea in vivo displayed a thicker epithelium in affected clones (green) compared to the neighboring cells (B). The corresponding cells were not thicker in the control trachea (A). (C-F) Trachea of *vvl-coin* larvae (C) *vvl-coin>Hop.CA* larvae (D-F) were transferred to PBS (D) or Schneider's medium (F) or fixed immediately in 4% paraformaldehyde (E). All of them showed a significant increase in cell volume. Scale bar: 100 µm in A-B; 20 µm in C-F. 50 specimens are used in each experiment.

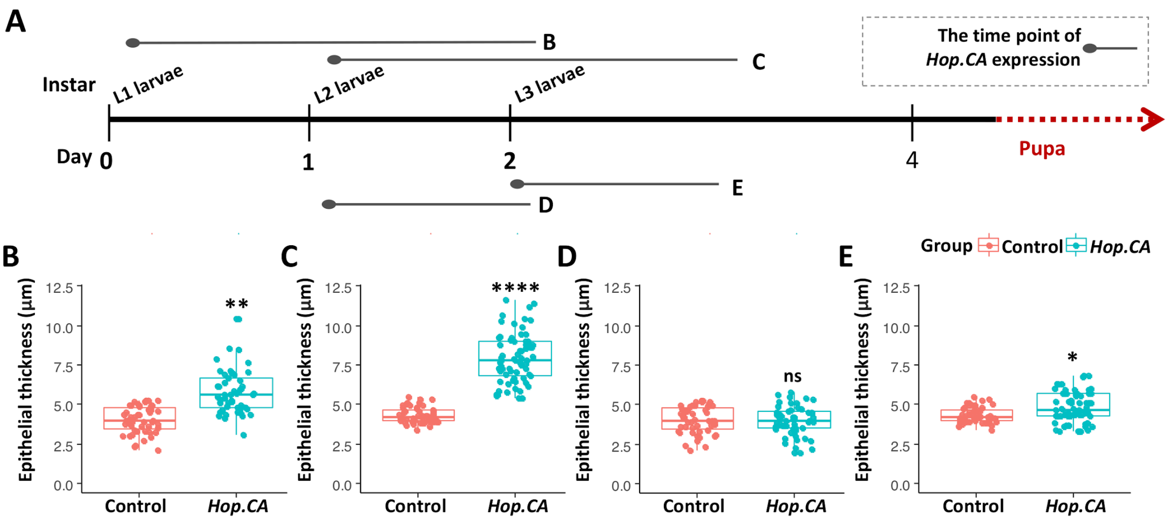


Figure S5 Time point and period of activation of *Hop.CA* expression by using the *btl.ts* driver. (A) Illustration of differing start and end points of *Hop.CA* expression by exposing them to the permissive temperature for one or two days in different larval stages. (B-E) Quantification of epithelial thickness of the DT8 region of control larvae (*btl.ts* driver line) and those experiencing ectopic manipulation (*btl.ts>Hop.CA* line). Each group contains 50 replicates in D-E. ns means not significant, * $p < 0.05$, ** $p < 0.01$, **** $p < 0.0001$ by Student's t-test.

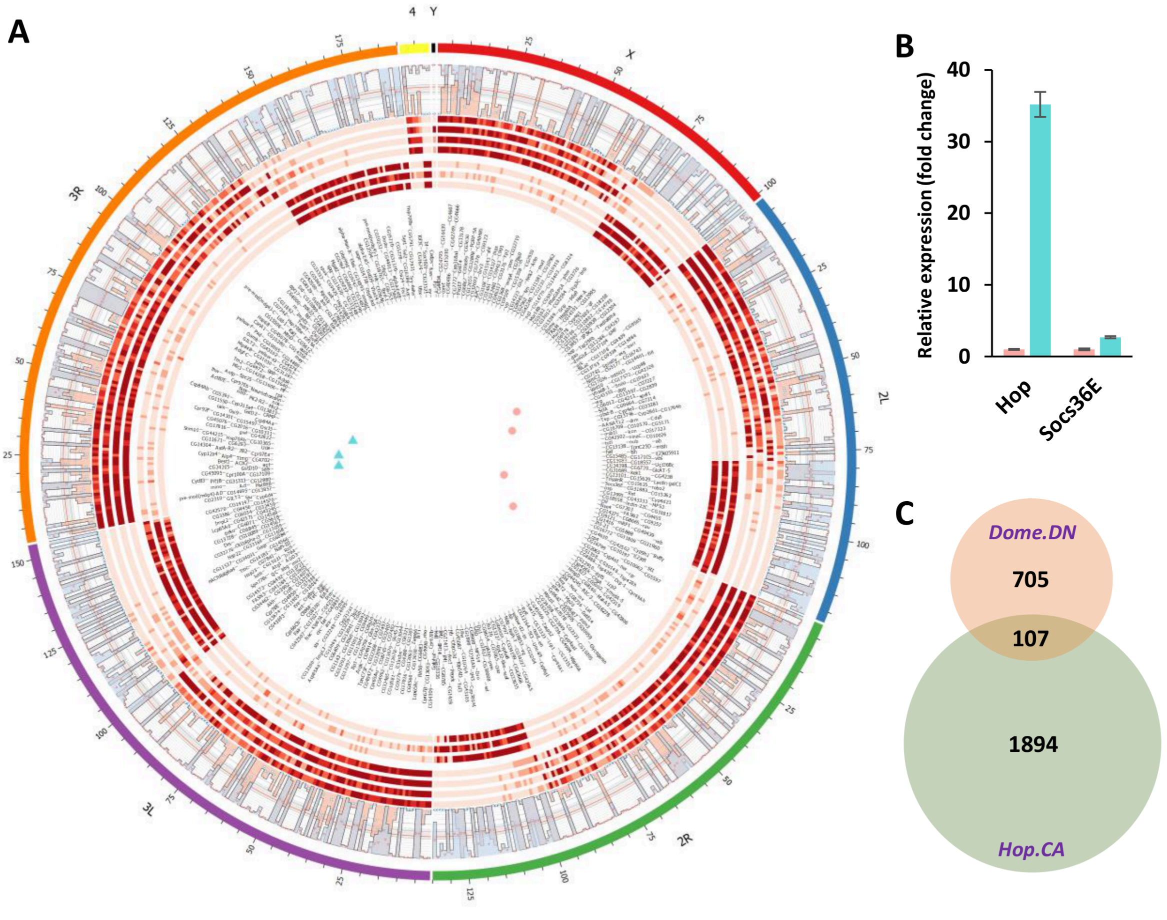

Figure S6: Changes at the transcript level in the airway epithelium of JAK/STAT mutants. Altered gene expression levels in the trachea caused by 16 hours persistent expression of *Hop.CA* driven by *btl.ts*. In total, 2004 genes were regulated significantly ($p < 0.05$), in which 1128 genes were down-regulated, 876 genes were up-regulated. (A) 707 genes (fold change > 2, $p < 0.01$) were used to visualize the differences of the transcript levels between *Hop.CA* expressed trachea and control trachea. From the outside to the inside, a histogram for expression mean values (red, control; blue, *Hop.CA*), a heatmap for controls, a heatmap for *Hop.CA* overexpression, gene names, and PCA analysis, are shown. According to transcriptome analysis of the tracheal epithelium 1128 genes ($p < 0.05$) were down-regulated, 876 genes ($p < 0.05$) were up-regulated. The changes in the gene expression total value were not significant in the ectopic expression trachea compared to the controls. (B) Relative expression levels of *Hop* and *Socs36E* in *Hop.CA* trachea compared to control trachea. (C) The inhibition of the JAK/STAT signaling by ectopic expressing *Dome.DN* for 18 hours affects the transcription of 812 genes, however, which only 102 genes were regulated in both types of airways; with ectopically activated and inhibited JAK/STAT signaling.

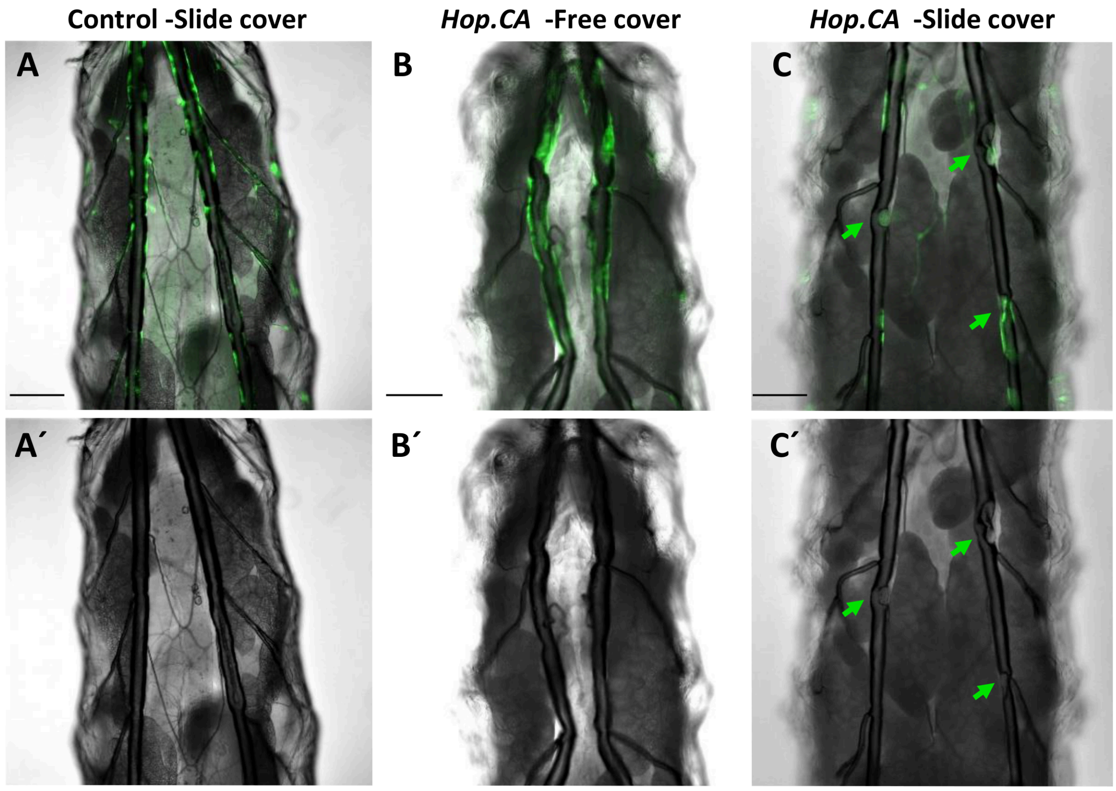


Figure S7 Micrographs of the trachea of *vvl-coin* larvae under different conditions. In A, the control is shown, in B, the undisturbed larvae of animals experiencing *Hop.CA* overexpression are displayed. (C) similar animals as in B, but the animals covered with a slide, inducing a certain degree of compression. The arrows show the collapsed sites in the tracheal tube. 30 larvae in each group were investigated in A-C. Scale bar: 200 µm.

### Supplementary tables

Table S1: Kegg pathway analysis of genes, whose expression is regulated in response to *Hop.CA* overexpression.

| Term | # regulated gene | #REF | corrective p-value |
|---|---|---|---|
| Protein processing in endoplasmic reticulum | 37 | 122 | 0.000196415 |
| Metabolism of xenobiotics by cytochrome P450 | 21 | 62 | 0.005458041 |
| Glutathione metabolism | 20 | 62 | 0.006987233 |
| Drug metabolism - cytochrome P450 | 20 | 62 | 0.006987233 |
| Metabolic pathways | 143 | 924 | 0.010283682 |
| Fatty acid metabolism | 14 | 42 | 0.033601606 |

Table S2: 32 regulated genes in GO9 and GO10.

| Name | Max group mean | Fold change | FDR p-value |
|---|---|---|---|
| CG4293 | 30.32889 | 1.492049 | 0.004753 |
| Cog7 | 15.97701 | 1.44363 | 0.012736 |
| deltaCOP | 103.7106 | 1.56551 | 0.00068 |
| CG9536 | 23.15629 | 1.351438 | 0.02827 |
| Sec22 | 72.30199 | 1.402676 | 0.031492 |
| sau | 77.49881 | 1.431877 | 0.01605 |
| epsilonCOP | 76.10362 | 1.399189 | 0.011535 |
| CG7456 | 23.9483 | 1.537314 | 0.001188 |
| Syx18 | 28.09394 | 1.355287 | 0.021462 |
| CG11857 | 79.55307 | 1.348644 | 0.047345 |
| zetaCOP | 96.3113 | 1.510022 | 0.003362 |
| CG31729 | 46.75703 | 1.881828 | 7.98E-07 |
| CG5946 | 43.55012 | -1.48145 | 0.006965 |
| CG7011 | 52.77113 | 1.986896 | 2.63E-08 |
| alphaCOP | 74.65062 | 1.44925 | 0.029706 |
| GABPI | 24.92601 | 1.662457 | 5.17E-05 |
| CG4293 | 30.32889 | 1.492049 | 0.004753 |
| p115 | 23.55302 | 1.500032 | 0.00275 |
| bai | 264.6093 | 1.394544 | 0.025263 |
| KdelR | 165.6429 | 1.434894 | 0.034105 |
| betaCOP | 95.96036 | 1.602758 | 0.001899 |
| Bet1 | 36.37198 | 1.509118 | 0.005981 |
| Sec24AB | 22.08689 | 1.427689 | 0.012235 |
| Tango1 | 26.83374 | 1.451327 | 0.021437 |
| Sec13 | 103.0673 | 1.556802 | 0.000783 |
| PAPLA1 | 18.71817 | 1.703538 | 5.78E-05 |
| Sec16 | 12.41264 | 1.385734 | 0.041951 |
| Sec23 | 122.68 | 1.544822 | 0.010929 |
| Sec31 | 49.0174 | 1.658867 | 0.000545 |
| loj | 137.7512 | 1.573849 | 0.000975 |
| Sec24CD | 53.79686 | 1.555751 | 0.002476 |
| eca | 159.2751 | 1.512845 | 0.003406 |
| ergic53 | 180.2582 | 1.749912 | 0.000928 |
| alphaCOP | 74.65062 | 1.44925 | 0.029706 |

Table S3: Transcription factor-binding site motifs enriched in 32 regulated genes in GO9 and GO10.

| Matrix ID | Matrix Name | P-value |
|---|---|---|
| MA0230.1 | lab | 0.00226225 |
| MA0244.1 | slbo | 0.00555053 |
| MA0446.1 | fkh | 0.00663537 |
| MA0186.1 | Dfd | 0.010245 |
| MA0165.1 | Abd-B | 0.0114577 |
| MA0170.1 | C15 | 0.0120483 |
| MA0013.1 | Br (var.4) | 0.0219955 |
| MA0203.1 | Scr | 0.0261687 |
| MA0206.1 | abd-A | 0.0264781 |
| MA0448.1 | H2.0 | 0.0340574 |
| MA0197.2 | nub | 0.0356034 |
| MA0174.1 | Dbx | 0.0369558 |
| MA0458.1 | slp1 | 0.0372846 |
| MA0221.1 | eve | 0.0412541 |
| MA0219.1 | ems | 0.0429901 |
| MA0225.1 | ftz | 0.0474096 |
| MA0012.1 | Br (var.3) | 0.0476299 |
